# Synthetic reaction-diffusion patterning in mammalian cells

**DOI:** 10.64898/2026.11.01.763437

**Authors:** Sheng Wang, Rongrong Du, James M. Linton, Michael B. Elowitz

**Author notes:** Wyss Institute for Biologically Inspired Engineering, Harvard University, Boston, MA, USA. Department of Biology, Stanford University, Stanford, CA, USA. These authors contributed equally to this work (Sheng Wang, Rongrong Du).

## Abstract

During development, periodic patterns arise through reaction–diffusion (RD) circuits involving local activation and long-range inhibition (LALI). Natural systems are complex, typically entangling regulation and transport in the same morphogens. It remains unclear what minimal circuits are sufficient for LALI patterning in mammalian cells, and whether they can be implemented in a modular fashion, allowing independent control of regulation and transport. Here, we designed synthetic LALI circuits using three natural and synthetic signaling systems—BMP, Activin, and the SNIPR pathway—and showed that they generate spatial patterns with distinct characteristic spatial scales. Model-guided design allowed flexible morphogen reassignment, with BMP4 acting as either short-range activator or long-range inhibitor. Gene expression noise degraded long-range order but preserved the dominant local spatial scale. Together, these results provide a modular, extensible synthetic framework for engineered patterning in mammalian cells.

## Introduction

Repetitive structures are pervasive across metazoan development, underlying teeth (*1*), limbs (*2*), fingerprints (*3*), hair follicles (*4*), and animal coat patterns (*5*). These patterns can arise from local interactions without centralized control. Alan Turing’s reaction–diffusion (RD) theory provides a foundational framework for periodic patterning, with local activation and long-range inhibition (LALI) as its core principle (*6*). In vivo, native patterning circuits are deeply entangled with developmental programs, making it difficult to isolate and study core pattern-forming modules.

Synthetic biology offers an alternative approach. Engineered cell–cell communication has long been used to program spatial patterns in bacteria(*7*), and recent work has extended synthetic RD engineering to intracellular dynamics in mammalian cells (*8*). For multicellular mammalian patterning, building minimal LALI circuits from defined molecular components provides a way to isolate core LALI circuits from growth, mechanics, and fate maturation, and offers a controlled setting to compare distinct circuit designs. Previous efforts focused on reconstituting the natural Nodal/Lefty system (*9*). In that case, the same diffusible molecules transmit spatial information and implement activation–inhibition logic, coupling signal transport to intracellular reaction and computation. Decoupling spatial communication from intracellular response logic would allow each layer to be independently engineered, tuned, and recombined, as demonstrated by bacterial quorum-sensing circuits that generate stochastic Turing patterns (*10*). Whether an analogous decoupled LALI architecture can drive self-organized patterning in mammalian cells remains unclear.

Here, we establish a synthetic LALI architecture in mammalian cells, in which extracellular signals transmit spatial information while activator or inhibitor roles are assigned by intracellular interfaces and regulatory modules. Building on modular synthetic receptor and cell–cell signaling platforms(*11–15*), we implemented this design in two circuits—BMP4/Activin and SNIPR (SyNthetic Intramembrane Proteolysis Receptors)/BMP4—each of which self-organizes RD-like periodic patterns in confluent monolayers. Critically, the same diffusible molecule, BMP4, can be wired through different intracellular interfaces to occupy opposite network roles: an activator-driving signal in one circuit and an inhibitor-driving signal in the other. Circuit role is therefore specified by interface wiring rather than molecular identity, directly demonstrating the modularity and portability of the architecture. Consistent with the differing effective diffusion ranges of their activators, the two circuits produced patterns with distinct characteristic pattern scales.

The decoupled design allowed us to develop a lattice-ODE model incorporating a representative level of cell-to-cell expression heterogeneity. This model recapitulated the key statistical features of the experimental patterns and showed that cell-to-cell expression variability alone is sufficient to preserve dominant spatial scale selection while eroding long-range positional order. Together, these results provide an engineerable framework for synthetic patterning, and show how core circuits can establish specific patterning lengthscales, even as additional refinement may be necessary to achieve precise ordering.

## Results

### A framework for modular design of synthetic reaction-diffusion circuits

We first defined a generic design grammar for LALI circuits (Fig. 1A). A minimal decoupled LALI circuit requires two communication modules with different effective spatial ranges and one intracellular computation module. The shorter-range communication signal locally promotes production of both communication signals, thereby implementing local positive feedback. The longer-range signal activates an intracellular inhibitory module that suppresses the local activation branch. This architecture preserves the essential LALI logic — local amplification coupled to broader suppression — while allowing the communication channels and regulatory logic to be selected and rewired independently.

**Fig. 1.**
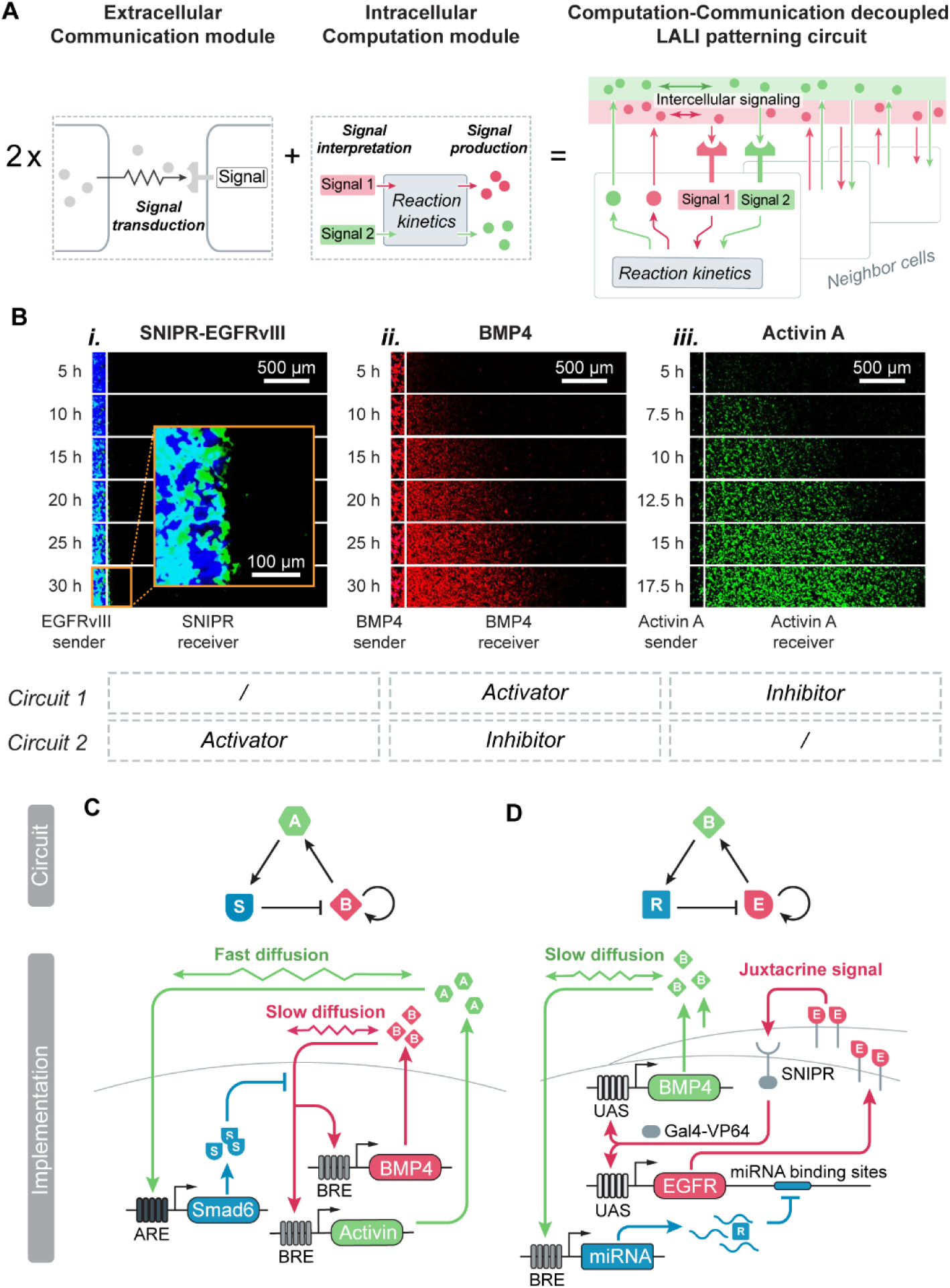
Design of communication–computation decoupled LALI circuits. (A) Two communication modules provide signals with different effective spatial ranges. The shorter-range signal promotes production of both communication signals, generating local positive feedback, whereas the longer-range signal activates an intracellular inhibitory module that suppresses the local activation branch. (B) Experimental validation of candidate communication modules using a one-dimensional sender–receiver assay. SNIPR–EGFRvIII signaling is restricted to direct cell–cell contact, BMP4 forms an intermediate-range gradient, and Activin forms a longer-range gradient, providing an ordered toolkit of communication channels with distinct spatial ranges. (C) Molecular implementation of the BMP4/Activin circuit. BMP4 signaling activates a BRE-driven module that promotes production of both BMP4 and Activin. Activin signaling activates an ARE-driven Smad6 module, which inhibits BMP signaling and closes the long-range inhibitory feedback loop. System: BMP4, Activin; Signaling Distance: Short, Long. (D) Molecular implementation of the SNIPR/BMP4 circuit. SNIPR activation drives production of both EGFRvIII and BMP4, forming a contact-dependent local activation branch. BMP4 signaling activates a BRE-driven synthetic microRNA that represses EGFRvIII, thereby inhibiting the activator branch. System: EGFRvIII-SNIPR, BMP4; Signaling Distance: Short, Long. Together, these designs establish a modular mammalian LALI framework in which communication range and intracellular regulatory logic can be recombined to generate distinct pattern-forming topologies.

To build this architecture in mammalian cells, we assembled and validated a toolkit of three communication channels spanning distinct effective spatial ranges: the synthetic juxtacrine SNIPR–EGFRvIII pathway, BMP4, and Activin (Fig. 1B and fig. S1). We quantified their spatial signaling ranges using quasi-one-dimensional sender–receiver time-lapse assays, in which localized sender cells produced each signal and receiver cells reported pathway activation across the surrounding field (Fig. 1B and figs. S1 and S2). SNIPR activation was restricted to cells in direct contact with EGFRvIII-expressing senders, giving it an operationally near-zero signaling range. BMP4 and Activin formed broader signaling gradients, with Activin spreading farther than BMP4 (Fig. 1B and fig. S2). Fitting the time-lapse gradient dynamics yielded effective signaling-spread coefficients (D_eff_) of 1.65 µm²/s for BMP4 and 10.03 µm²/s for Activin, consistent with their relative spatial ranges (fig. S2). These reporter-derived coefficients integrate transport and downstream response kinetics and are not direct measurements of molecular diffusivity. Thus, these three channels provide an ordered set of communication modules ranging from contact-dependent signaling to longer-range diffusible morphogen signaling (fig. S1). Dual-reporter perturbation assays further showed that BMP4/BRE and Activin/ARE responses were largely separable over the concentration range used for circuit design (fig. S3).

The two circuits implement the same abstract LALI logic through different molecular interfaces (Fig. 1, C and D), while deliberately placing BMP4 in opposite network roles — a direct test of whether circuit function is specified by intracellular wiring rather than molecular identity. Before assembling the full feedback circuits, we validated the corresponding open-loop regulatory modules, confirming that the local activation branches drove signal production and that the long-range-signal-induced inhibitory modules could repress those activation branches (fig. S4). In Circuit 1, BMP4 signaling activates a BRE (BMP4-responsive element)-driven module that promotes production of both BMP4 and Activin, forming the local amplification branch. Activin signaling activates an ARE (Activin-responsive element)-driven Smad6 module, which inhibits BMP4 signaling(*16*, *17*) and thereby provides delayed lateral inhibition (Fig. 1C). In Circuit 2, SNIPR activation drives production of both the EGFRvIII ligand and BMP4, forming a contact-dependent local activation branch(*18*, *19*). BMP4 signaling then activates a BRE-driven synthetic microRNA(*20*) that represses EGFRvIII, closing the inhibitory feedback loop onto the activator branch (Fig. 1D).

Across both circuit implementations, mCherry and mCitrine reported the activities of the activator and long-range inhibitory branches, respectively. Each reporter was expressed from the same bicistronic transcript as its corresponding circuit component as an H2B-fused, trimethoprim (TMP)-regulated DHFR destabilization-domain reporter (*21*). Addition of TMP at the start of the patterning assay stabilized newly synthesized reporters while minimizing fluorescence accumulated before pattern initiation (fig. S5).

Together, these designs establish a modular mammalian LALI framework in which extracellular communication channels define spatial range, intracellular computation modules define regulatory logic, and recombining the two layers generates alternative pattern-forming topologies. Because the activator signal changes from diffusible BMP4 in Circuit 1 to contact-dependent SNIPR signaling in Circuit 2, the framework predicts that the two circuits should generate patterns with distinct characteristic pattern scales. In the following sections, we test whether these designed circuits can self-organize spatial patterns and whether their behavior follows the expected LALI logic.

### Synthetic circuits produce spatial patterns in cell culture

Random genomic integration is typically viewed as an unwanted source of clone-to-clone variability in stable cell-line construction. Here, however, this variability provided a practical means of sampling circuit parameter space. Synthetic RD patterning is expected only within a restricted range of relative activation and inhibition strengths—the Turing-unstable regime. We therefore introduced the BRE-driven BMP4/Activin and ARE-driven Smad6 modules sequentially by piggyBac transposition and isolated approximately 300 full-circuit monoclonal lines after the second integration. To identify clones in which both circuit branches were functional, we first measured the two reporter outputs and excluded strongly imbalanced clones, such as those with high mCherry but low mCitrine, or vice versa. Importantly, this pre-screen was based on reporter detectability and balance rather than the presence or quality of spatial patterns. Thirteen clones meeting these predefined criteria were subsequently evaluated under standardized patterning conditions (fig. S6, Methods). Thus, the monoclonal lines used for quantitative analyses represent independently isolated circuit implementations within the relevant expression regime rather than clones selected for unusually strong patterning. The analyses below focus primarily on clone 22J, with key findings assessed across additional independently isolated clones.

With clones in hand, we performed pattern formation assays. Briefly, before patterning, we individually cultured each monoclonal line, adding the BMP4 inhibitor Noggin(*22*) to maintain cells in the “off” state, and avoid pre-patterning prior to the experiment. Then, we replated these cells and allowed them to settle with Noggin protection. After cells attached to the plate surface, we proceeded to the pattern formation assay. These procedures placed cells of identical genotype in a common suppressed, confluent starting condition before pattern initiation, with limited cell division during patterning.

After washing away Noggin, we initiated the experiment in a pattern-formation medium containing TMP (for fluorescent protein stabilization), Palbociclib (to inhibit cell division), and specific concentrations of DMH1 (BMP4 signal inhibitor) and A83-01 (Activin signal inhibitor) to tune circuit parameters (*23*). Full-field time-lapse imaging showed that clone 22J progressively developed spatially heterogeneous BRE- and ARE-reporter activity over 60 h (Fig. 2A, left and middle; Movie S1). By contrast, saturating A83-01 blocked Activin signaling and converted the circuit to a self-activation-only state, producing broad, nearly uniform BRE activation (Fig. 2A, right; Movie S2).

**Fig. 2.**
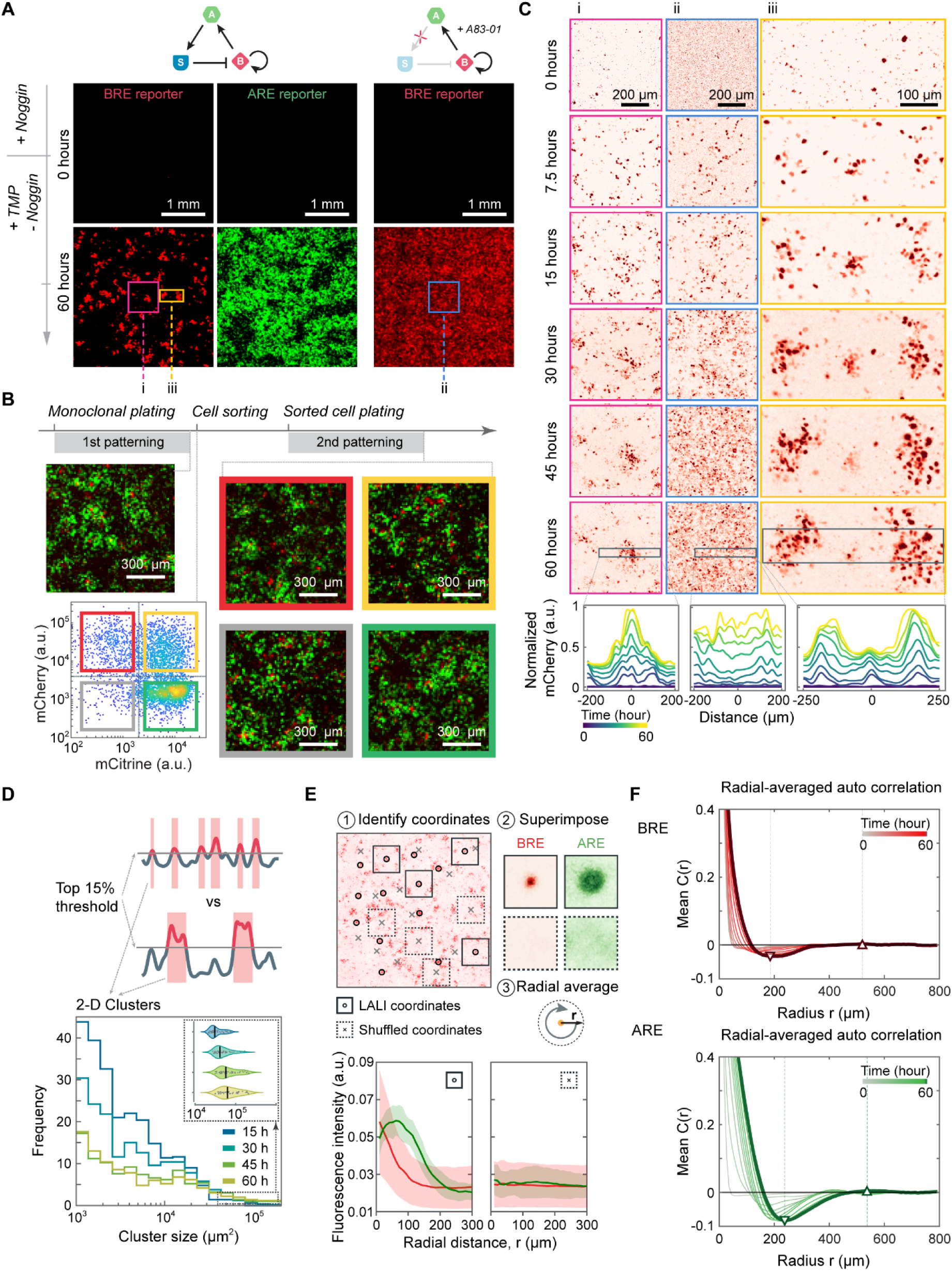
Synthetic BMP-Activin circuit generates spatial patterns. (A) Under patterning conditions, monoclonal Circuit 1 cells developed spatially heterogeneous reporter patterns from an initially suppressed state. Localized regions of high BRE reporter activity marked BMP4-driven activation domains, whereas ARE reporter activity reported the Activin/Smad6 inhibitory branch. In contrast, blocking Activin signaling with saturating A83-01 converted the circuit to a BMP4 self-activation-only control and produced more spatially uniform activation. (B) Sorted patterning cells reset their reporter states and re-formed spatial patterns after replating, indicating that the high- and low-reporter states are not stably inherited cell-autonomous states. (C) Time-lapse imaging captured the emergence and refinement of active domains. (i) Under patterning conditions, initially sparse BRE-positive cells seeded local active domains that expanded and stabilized over time. (ii) When Activin signaling was blocked, activation spread more broadly across the field, consistent with loss of the long-range inhibitory arm. (iii) Neighboring active domains were separated by regions of reduced activation, consistent with competition mediated by lateral inhibition. (D) Domain-size analysis from five independent movies showed progressive domain competition during Circuit 1 pattern formation. The top 15% brightest BRE-reporter pixels were segmented into connected clusters at each time point. Across movies, small active domains decreased over time, whereas the largest domains persisted or increased in area. The self-activation-only control did not show the same systematic change (fig. S7A). The trend was robust across smoothing widths, brightness thresholds, and largest-domain cutoffs used for quantification (fig. S7, B to D). (E) Event-based superposition analysis from five independent movies revealed a LALI-like spatial signature. Active-domain centers were identified from the BRE reporter channel and used to align local BRE and ARE reporter fields. Radial averaging showed local BRE enrichment near active-domain centers and broader ARE enrichment over surrounding regions. Coordinate-shuffled controls abolished this structure. (F) Radially averaged autocorrelation analysis from five independent movies showed a reproducible short-range negative correlation and a weak off-center positive peak, indicating local spatial organization with limited long-range order. Individual-movie autocorrelation curves and FFT-based pattern-spacing estimates are shown in fig. S8.

The heterogeneous reporter states were not stably inherited by individual cells. To test this, we sorted cells from distinct reporter-defined states after three days of patterning and replated them in fresh patterning medium. Rather than maintaining their original fluorescence states, the sorted populations reset and re-established heterogeneous patterns (Fig. 2B). Thus, the reporter states observed in the patterned culture depend on the local patterning environment rather than reflecting fixed cell-intrinsic states.

Zoomed-in time-lapse imaging resolved the local dynamics of domain formation under patterning and self-activation-only conditions (Fig. 2C, i and ii; Movie S3). Under patterning conditions, initially homogeneous cells sporadically activated the BRE promoter, producing scattered mCherry-positive cells during the first 7.5 h. These cells subsequently activated neighboring cells and seeded expanding clusters between 7.5 and 15 h. From 15 to 45 h, some clusters grew larger and brighter while others diminished. By 45–60 h, the remaining clusters approached a slowly changing configuration with characteristic sizes and spacing (Fig. 2C, i; Movie S3, left). In the self-activation-only control, in which the Activin arm was inhibited, sporadic activation was also evident by 7.5 h. However, activation subsequently spread across most of the field, producing a more spatially uniform state by 60 h (Fig. 2C, ii; Movie S3, right).

During pattern refinement, neighboring active domains appeared to compete as their inhibitory zones began to overlap (Fig. 2C, iii; Movie S4). Early in the movie, individual cells or small clusters became active at dispersed locations. As these domains expanded, some regressed whereas others persisted and continued to grow. By the end of imaging, a subset of spatially separated domains remained, producing an organized pattern.

To confirm that these dynamics are general rather than rare events, we quantified domain numbers and sizes using an 85th-percentile brightness threshold (Fig. 2D, five independent movies, n = 5). We assessed the size distribution of bright-pixel clusters, finding that many small clusters at early time points give way to fewer but larger clusters at later stages. We also plotted the sizes of the top 10 largest clusters at each time point, which further underscores the emergence of larger domains over time (Fig. 2D, inset). In the self-activation-only control, however, the size distribution remained evenly distributed (fig. S7A, four independent movies, n = 4). This conclusion was robust to the choice of Gaussian smoothing width, brightness-threshold percentile, and the number of largest domains used for area quantification (fig. S7, B to D).

We next employed an event-based superimposition strategy to visualize local interactions (Fig. 2E, five independent movies, n = 5). We identified the positions of all active domains (examples marked as circles) in the mCherry channel using a Laplacian of Gaussian (LoG) filter, which detects peaks in the smoothed image. Sub-images (n = 44 from 5 movies) were then extracted from both mCherry and mCitrine channels at these coordinates and superimposed, aligning domain centers. Radially averaging the resulting composite revealed that the self-activation signal (mCherry) is narrowly distributed, whereas the lateral inhibition signal (mCitrine) was spread more broadly, consistent with LALI. A null model generated by shuffling the coordinates (marked as crosses, n = 88 from the same 5 movies) showed uniform distributions in both channels, confirming the non-random nature of the observed pattern.

Finally, we quantified the spatial organization of the patterns using autocorrelation analysis after background subtraction and correction for uneven illumination (Fig. 2F; five independent movies, n=5). Radially averaged autocorrelation curves from both the mCherry and mCitrine channels showed a reproducible short-range negative correlation followed by a weak first off-center positive peak, indicating local spatial organization but limited long-range order. Complementary radial Fast Fourier Transform (FFT) analysis across the five independent movies yielded a characteristic pattern spacing of 546.8±54.9 μm using the FFT-centroid estimator. Individual-movie autocorrelation and FFT profiles, together with spacing estimates from multiple methods, are shown in fig. S8.

Together, these analyses confirm that the synthetic reaction–diffusion circuit can drive pattern formation through local activation with long-range inhibition.

### Cell-to-cell promoter heterogeneity preserves local spatial scale while rapidly eroding long-range order

Autocorrelation analysis of the Circuit 1 patterns revealed a characteristic local spacing identified by both FFT and autocorrelation analyses, together with rapid spatial correlation decay (Fig. 2F). This indicated that local periodicity was established without precise long-range positional order. Whether this loss of long-range order reflected experimental conditions or an intrinsic sensitivity of the LALI core to cell-to-cell heterogeneity could not be determined from the experimental data alone. Cell-to-cell variation in kinetic parameters is known to reshape reaction–diffusion patterns by perturbing local reaction rates and altering the regions of parameter space that support pattern formation (*24*, *25*). We therefore asked whether the limited spatial coherence of the experimental patterns could be explained by cell-to-cell expression heterogeneity.

We developed a lattice-based ODE model of Circuit 1 in which each node represents one cell, the circuit equations are integrated at every node, and diffusible species couple neighboring nodes (Fig. 3A; Methods). This cellular representation allowed us to incorporate expression differences directly at the single-cell level while retaining a framework that can also accommodate the contact-dependent SNIPR interactions in Circuit 2. To represent cell-to-cell expression heterogeneity, each cell was assigned a single mean-corrected lognormal multiplier, parameterized by σ, which scaled the BMP4, Activin, and Smad6 production rates within that cell (Fig. 3A). Here, σ controls the cell-to-cell variation of all production rates on the log scale. The mean correction ensures that changing σ alters the spread of the parameter distribution without systematically shifting the population-averaged production rates.

**Fig. 3.**
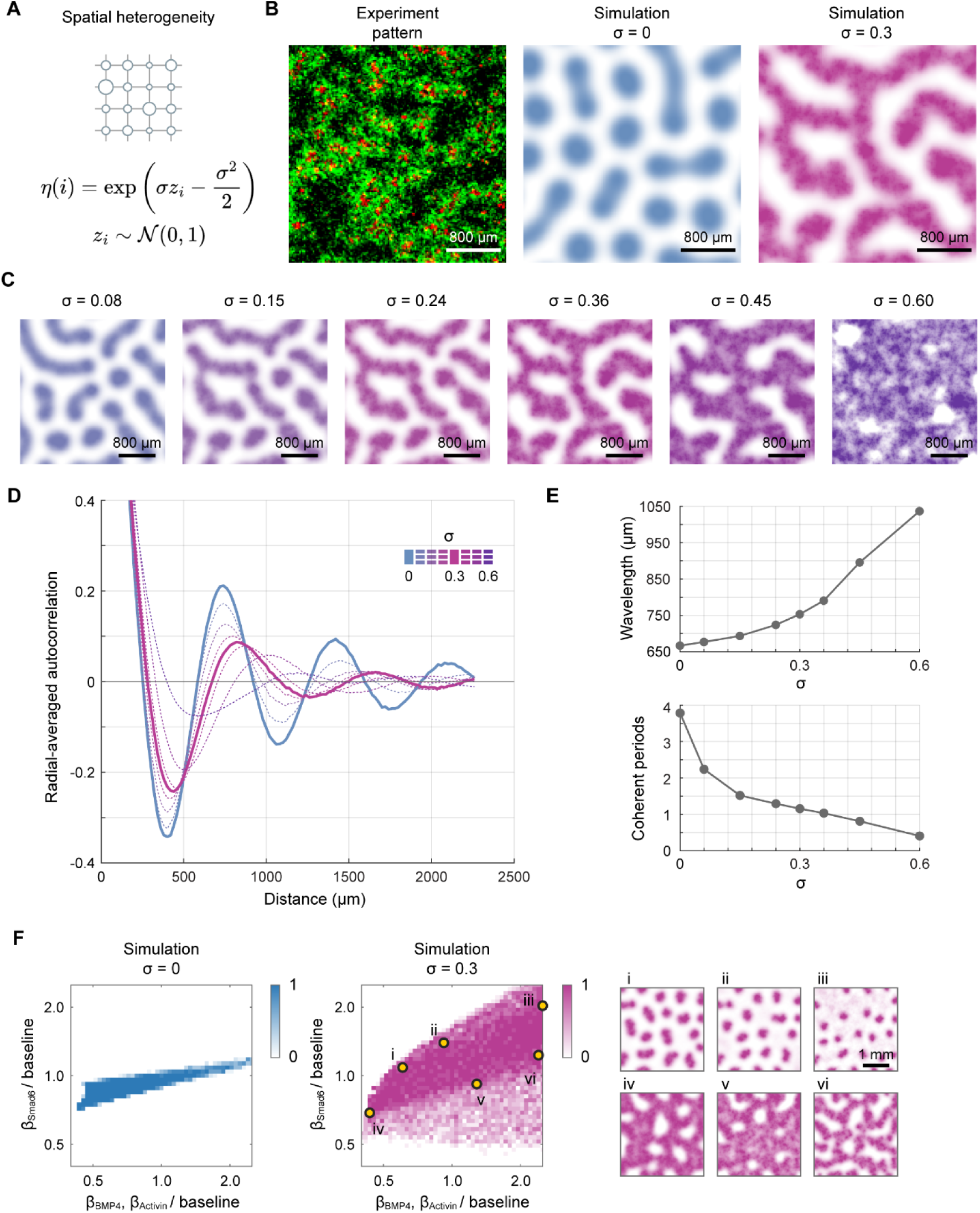
Cell-to-cell expression heterogeneity erodes long-range order and expands the pattern-forming region. (**A**) Each cell is assigned a mean-corrected lognormal multiplier, η(i), parameterized by σ, that scales its BMP4, Activin, and Smad6 production rates. (**B**) Experimental Circuit 1 pattern (ARE–mCitrine, green; BRE–mCherry, red) compared with simulations at σ = 0 and σ = 0.3. Heterogeneity produces diffuse, irregular domains resembling the experiment. (**C**) Representative patterns across the indicated σ values. (**D**) Radially averaged autocorrelation, C(r), across the heterogeneity titration. Increasing σ attenuates successive off-center peaks and modestly shifts them toward larger distances. (**E**) Wavelength increases only modestly, whereas coherent periods (ξ/λ) decrease sharply at low σ and approach one at higher σ. (**F**) Pattern-formation probability from 10 replicate simulations across joint BMP4/Activin and Smad6 production scales normalized to baseline. Patterns occupy a narrow diagonal band at σ = 0 but a substantially broader region at σ = 0.3. Insets i–vi show representative patterns.

The spatially homogeneous model (σ = 0) generated labyrinthine patterns, but the domains were overly regular, with smooth boundaries and a degree of spatial coherence not observed experimentally (Fig. 3B). Introducing a representative level of cell-to-cell heterogeneity (σ = 0.3) (*26*, *27*) roughened the domain boundaries and produced a more diffuse local texture, yielding a pattern morphology that more closely resembled the experimentally observed finite-coherence patterns. A single source of cell-to-cell expression heterogeneity was therefore sufficient to convert the excessively regular deterministic pattern into a spatial organization more representative of the experiment.

To determine how pattern morphology depended on heterogeneity magnitude, we increased σ from 0.08 to 0.60 (Fig. 3C). As σ increased, domain boundaries became progressively less regular, positional alignment between distant regions weakened, and the pattern coarsened, with a corresponding increase in characteristic spacing. Recognizable spatial patterns persisted even at the higher heterogeneity levels examined, but their organization became increasingly local rather than remaining coherent across multiple periods.

The radially averaged autocorrelation function, C(r), quantified these changes (Fig. 3D). At low heterogeneity, C(r) displayed multiple persistent off-center peaks, indicating periodic order extending across several wavelengths. As σ increased, the successive autocorrelation peaks decayed rapidly, demonstrating a progressive loss of phase relationships between distant regions of the pattern. At the same time, the peaks shifted toward larger spatial distances, consistent with the increase in pattern scale observed in the simulated fields.

Wavelength and spatial coherence showed markedly different sensitivities to heterogeneity (Fig. 3E). As σ increased from 0 to 0.60, the pattern wavelength showed only a modest increase across the range examined. In contrast, the number of coherent periods, quantified as the ratio of the correlation length to the wavelength, ξ/λ, decreased sharply at low heterogeneity. It fell from nearly four periods at σ = 0 to approximately two periods at σ = 0.08, and then approached one period at higher σ. Long-range order was therefore substantially more sensitive to cell-to-cell heterogeneity than pattern wavelength, partially decoupling local length-scale selection from long-range positional order.

The same heterogeneity also substantially altered the range of circuit parameters that supported pattern formation. We scanned the relative strengths of the BMP4 and Activin production rates against that of Smad6 and compared pattern probabilities at σ = 0 and σ = 0.3 (Fig. 3F). In the spatially homogeneous model, pattern formation was confined to a narrow diagonal band, whereas most other parameter combinations converged to spatially homogeneous states. Introducing σ = 0.3 markedly expanded the pattern-forming region and increased pattern probability across multiple parameter combinations that did not form patterns in the homogeneous model. Representative parameter sets i–vi produced diverse spatial structures, ranging from isolated spot-like domains to connected labyrinthine patterns.

Cell-to-cell expression heterogeneity therefore had a dual effect on LALI patterning. At a given set of circuit parameters, increasing heterogeneity enlarged the characteristic wavelength and sharply reduced long-range spatial coherence. Across the broader parameter space, however, heterogeneity expanded the range of expression states capable of supporting pattern formation. Cell-to-cell expression variability should consequently be treated not merely as an experimental imperfection, but as a system-level property that jointly determines pattern scale, long-range order, and patternability.

### Circuit parameters and boundary conditions allow predictable modulation of pattern formation

In general, patterning can depend on circuit parameters and boundary conditions. To probe how circuit parameters set pattern morphology, we mapped inhibition of BRE promoter onto an increase in the activator half-max constant K_B_ and inhibition of ARE promoter onto an increase in the inhibitor half-max constant K_A_ in the lattice ODE model, and performed a 3×3 parameter scan around the calibrated point (Fig. 4A). The scan recovered three signatures of a Turing-type instability(*28–31*). First, when K_B_ and K_A_ are varied synchronously, pattern morphology is essentially unchanged, consistent with the dependence of Turing pattern selection on relative reaction parameters and nonlinear response structure. Second, raising K_B_ while lowering K_A_ (BMP signal inhibition-dominant regime) shifts the field from worm-like to discrete dots. Third, lowering K_B_ while raising K_A_ (Activin signal inhibition-dominant regime) merges active domains into a sponge-like morphology. Notably, in the deterministic scan (fig. S9) two opposing corners of parameter space collapse to homogeneous fixed points, whereas the noise-added scan recovers patterns in both corners—indicating that promoter noise expands the patternable region of parameter space on either side of the Turing-unstable band.

**Fig. 4.**
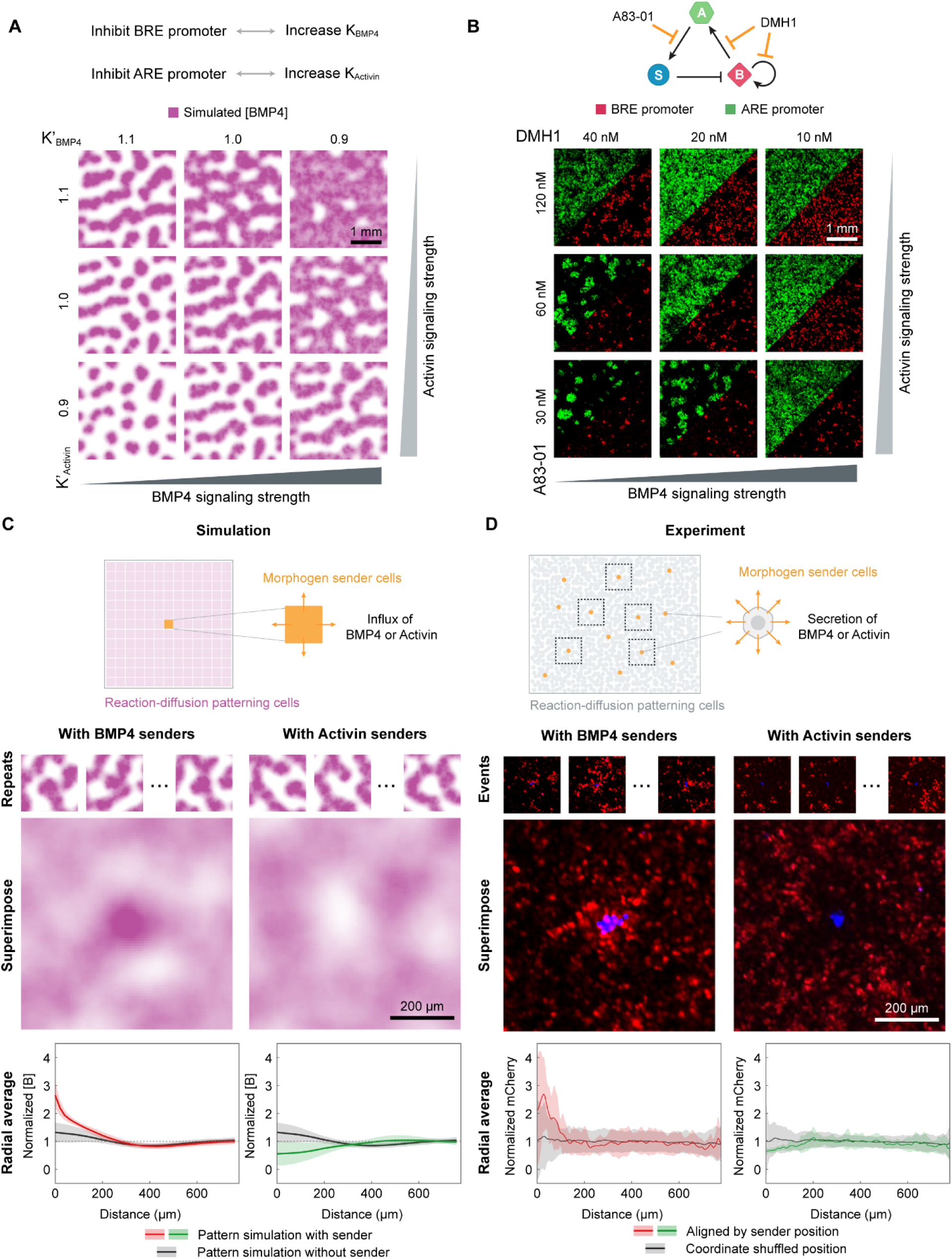
Predictable modulation of pattern formation. (A) A lattice-based reaction–diffusion ODE simulation predicts morphological transitions in response to modulated BMP4 and Activin signaling intensities. (B) Titration of two selective kinase inhibitors, DMH1 and A83-01, induces systematic changes in emergent spatial patterns. Enhancing BMP4 signaling or reducing Activin signaling increases both the number and size of active domains, shifting patterns from sparse dots to sponge-like structures. (C) Simulations mimicking BMP4 or Activin senders by modifying the central lattice node show active domain redistribution. (D) Co-culturing patterning cells with sparse morphogen senders reveals boundary-dependent pattern adaptation. BMP4 influx from the boundary promotes the formation of nearby active domains, while Activin influx repels them.

To experimentally test these predictions, we varied DMH1 and A83-01 concentrations across a 3×3 matrix (Fig. 4B). Decreasing DMH1 increased BMP signaling, whereas decreasing A83-01 increased Activin signaling. Across this matrix, the reporter patterns changed qualitatively in the direction predicted by the model. Conditions with weaker BMP activation and stronger Activin signaling, corresponding to high DMH1 and low A83-01, produced more spatially separated reporter-positive domains. In contrast, stronger BMP activation combined with weaker Activin signaling, corresponding to low DMH1 and high A83-01, produced more interconnected patterns. The BRE/mCherry and ARE/mCitrine channels changed coherently across this perturbation matrix, consistent with modulation of the coupled activation–inhibition circuit. Although the experimental patterns retained greater cell-scale texture than the simulations, their overall morphological transitions were qualitatively consistent with model predictions. Time-lapse imaging of an expanded independent titration showed the progressive emergence of the same condition-dependent morphological transitions across a finer DMH1/A83-01 matrix (fig. S10; Movie S5).

Reaction–diffusion patterns are sensitive to boundary conditions, which we leveraged as a complementary strategy in our perturbation studies. Previous work used recombinant morphogen protein-coated beads to introduce new boundaries through creating local morphogen influxes(*4*). To investigate whether boundary conditions re-direct the local pattern formation, we introduced point sources of either the short-range activator (BMP4) or the long-range inhibitor (Activin) and examined how they reshape the surrounding pattern.

We first simulated this perturbation in the lattice ODE model. “Sender” nodes with constant production of either BMP4 or Activin were introduced at random positions into otherwise standard Circuit 1 simulations, and the local field around each sender was superimposed over many realizations to extract a statistical radial response (Fig. 4C). The simulations predicted opposite effects of the two sender types. Surrounding BMP4 senders, the radial BRE response was sharply elevated—peak amplitude ∼3× the field mean—decaying to baseline within ∼200–300 µm, consistent with BMP4 acting as the local activator. Activin senders, by contrast, produced a decrease in the BRE response at the sender position that recovered to baseline over a similar length scale, consistent with the lateral-inhibition cascade Activin → Smad6 ⊣ BRE.

We then tested these predictions experimentally. We co-cultured patterning cells with sparse cells constitutively secreting either BMP4 or Activin, ran the patterning assay, and after pattern stabilization aligned all sender positions across many fields and superimposed the surrounding active-domain distributions (Fig. 4D). The resulting radial profiles closely matched the simulations: BMP4 senders locally attracted active domains, whereas Activin senders locally repelled them. A null model in which sender coordinates were shuffled produced flat radial profiles in both conditions, confirming that the observed responses were sender-driven rather than analysis artifacts.

Together, these experimental and simulation results provide strong evidence that our synthetic reaction–diffusion circuit functions as designed. By perturbing circuit parameters and boundary conditions, we observed predictable and reproducible changes in spatial patterns. The consistency between experimental data and simulations further validates the underlying design principles, confirming that the synthetic circuit exhibits hallmark behaviors of reaction–diffusion systems, including sensitivity to parameter tuning and boundary effects. These findings collectively demonstrate the robustness, responsiveness, and functional integrity of the engineered patterning system.

### The SNIPR/BMP4 circuit spontaneously and predictably develops spatial patterns with shorter length scales

To test the generality of our modular design framework, we constructed and characterized a second reaction–diffusion circuit in which SNIPR–EGFRvIII juxtacrine signaling serves as the short-range activator and diffusible BMP4 serves as the long-range inhibitor (Fig. 1D). Because SNIPR activation requires direct cell-cell contact, its effective spatial range is expected to be substantially shorter than that of the diffusible BMP4 activator in Circuit 1. This design therefore predicts that Circuit 2 should generate patterns with a shorter characteristic spatial scale(*32–34*).

Using the same clone-selection strategy established for Circuit 1, we identified SNIPR/BMP4 monoclonal lines capable of pattern formation and performed time-lapse imaging under patterning conditions (Fig. 5A; Movie S6). As with the BMP4/Activin circuit, initially homogeneous cells progressively developed spatially heterogeneous reporter activity over 60 hours. EGFRvIII/SNIPR juxtacrine signaling reporter activity (red) emerged as localized active domains, whereas BMP4/BRE reporter activity (green) extended across the surrounding field, consistent with BMP4 functioning as the longer-range inhibitory signal in this circuit. Quantification of the same movies showed a time-dependent redistribution of active-domain sizes as the pattern developed (fig. S11D). Event-based superimposition centered on active SNIPR/UAS domains further revealed reproducible local organization of the reporter signals, whereas shuffled coordinates produced approximately flat radial profiles (fig. S11E). Together, these analyses show that Circuit 2 generates non-random, locally organized spatial domains rather than isolated fluorescence fluctuations.

**Fig. 5.**
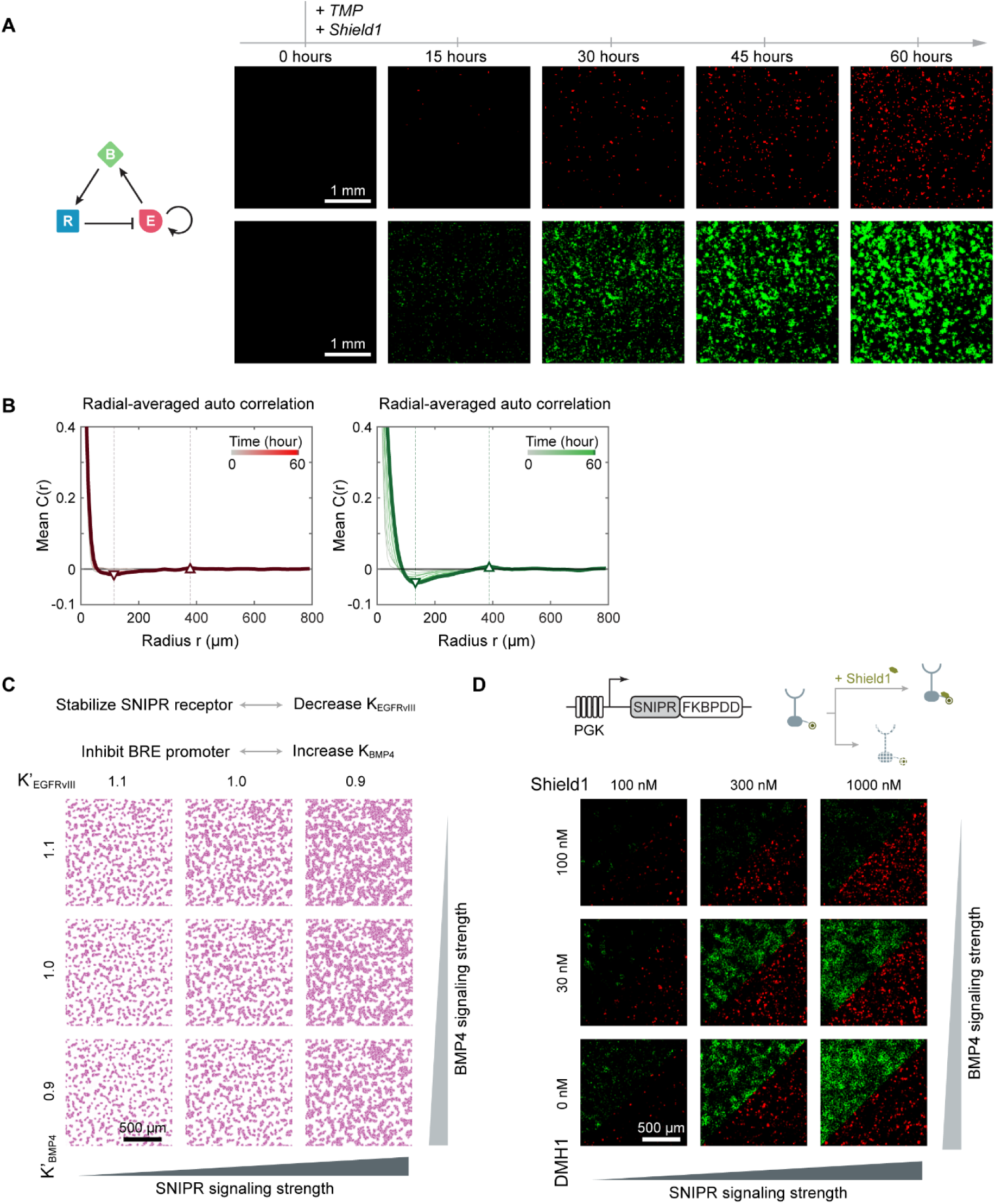
The SNIPR/BMP4 circuit generates tunable spatial patterns with shorter length scales. (A) Time-lapse imaging of SNIPR/BMP4 monoclonal cells under patterning conditions at 0, 15, 30, 45 and 60 hours. Initially homogeneous cells progressively develop spatially heterogeneous domains over 60 hours. SNIPR signal (red, top) and BMP4 signal (green, bottom) (B) Radial-averaged autocorrelation analysis of the SNIPR (red, left) and BMP4 (green, right) reporter channels. Both channels converge to a stable profile over the course of the experiment, revealing a characteristic domain spacing of approximately 319.3 ±15.3 µm. (C) The same lattice-based modeling framework used for Circuit 1, adapted to the hybrid juxtacrine–paracrine architecture of the SNIPR/BMP4 circuit, predicts systematic changes in domain density and connectivity as SNIPR stabilization and BMP4 signaling strength are varied. (D) Titration of Shield1, which stabilizes SNIPR, and DMH1, which attenuates BMP4 signaling, produces systematic and predictable changes in pattern density and morphology, which agree with simulation.

We next quantified the characteristic spatial scale of Circuit 2 patterns (Fig. 5B and fig. S11, A to C). After background and uneven-illumination correction, radially averaged autocorrelation curves from four independent movies showed a reproducible short-range negative correlation followed by a weak off-center positive peak, indicating local spatial organization but limited long-range order (fig. S11B). Radially averaged two-dimensional FFT spectra from the same movies showed dominant nonzero-frequency peaks corresponding to the characteristic pattern scale (fig. S11A). Because the off-center autocorrelation peaks were weak, we used the centroid of the dominant FFT peak as the primary spacing estimator and compared it with three alternative FFT- and autocorrelation-based estimates (fig. S11C). The FFT-centroid analysis yielded a characteristic spacing of 319.3 ±15.3 µm for Circuit 2. Using the same estimator, Circuit 1 showed a spacing of 546.8± 54.9 µm (fig. S8C), making the Circuit 1 spatial scale approximately 1.7-fold larger than that of Circuit 2. The alternative estimators produced the same qualitative conclusion (figs. S8C and S11C). This difference is consistent with the design prediction that replacing a diffusible BMP4 activator with contact-dependent SNIPR signaling compresses the characteristic pattern scale.

To predict how circuit parameters shape Circuit 2 pattern morphology, we performed a parameter scan in the lattice ODE model (Fig. 5C). Similar to Circuit 1, we mapped stabilization of SNIPR onto a decrease in the activator half-max constant K’_EGFRvIII_ and inhibition of BRE promoter onto an increase in the inhibitor half-max constant K’_BMP4_. The model predicted that Circuit 2 was more sensitive to SNIPR signaling strength than to BMP4 signaling strength: increasing SNIPR signaling produced more frequent and larger active domains, whereas changing BMP4 signaling over the same relative range had a weaker effect on pattern morphology. This asymmetry suggests that SNIPR receptor abundance is the dominant determinant of Circuit 2 pattern density, with BMP4 signal attenuation playing a secondary role. The corresponding noise-free simulations are shown in fig. S12; however, as observed for Circuit 1, simulations incorporating noise more closely recapitulated the patterns observed experimentally.

We then experimentally tested this prediction by titrating Shield1 and DMH1 (Fig. 5D). Shield1 strengthens the juxtacrine activation arm through stabilizing the FKBP destabilization domain-fused SNIPR receptor, thereby increasing receptor abundance and juxtacrine activation. DMH1 reduces BMP4 signaling, thereby weakening BRE-driven microRNA repression of the EGFRvIII/SNIPR activation branch. Across a matrix of Shield1 and DMH1 concentrations, Circuit 2 showed systematic and predictable changes in pattern density and morphology. At low Shield1 and low DMH1, active domains were sparse and well separated. Increasing Shield1 enhances SNIPR activation, increasing active-domain number and area, with this effect approaching saturation at higher concentrations. Increasing DMH1 also promoted active-domain expansion, consistent with reduced BMP4/BRE-driven microRNA repression and release of the EGFRvIII/SNIPR activation branch. Time-lapse imaging of an expanded Shield1/DMH1 titration matrix showed the progressive emergence of condition-dependent differences in active-domain density and morphology (fig. S13; Movie S7).

Together, these results show that the modular LALI framework can access distinct patterning regimes by substituting signaling modules with different spatial ranges while preserving the same core patterning logic. Importantly, BMP4 takes on opposite functional roles in the two circuits: in Circuit 1, BMP4 acts as the short-range activator, whereas in Circuit 2, BMP4 acts as the long-range inhibitor. The molecular identity of BMP4 is unchanged; what differs is its input-output wiring and downstream regulatory logic. Thus, activator or inhibitor identity is not a fixed molecular property of a morphogen, but an assignable functional role specified by circuit topology and intracellular wiring.

## Discussion

These results demonstrate that it is possible to design synthetic LALI circuits that generate self-organized spatial patterns in mammalian cell culture (Fig. 2; Fig.5). Our decoupled design enables independent selection of extracellular signaling modules, which determine how signals propagate between cells, and intracellular wiring, which determines how cells interpret those signals and generate subsequent signaling outputs (Fig. 1A). Using this principle, we constructed two distinct molecular implementations of LALI from overlapping sets of components (Fig. 1, C and D). The wavelength difference between the circuits is consistent with the shorter effective range of the local activator in Circuit 2, and with model predictions (Fig. 2F; Fig. 5B).

These circuits use BMP4 in opposite roles. It acts as the short-range activator in Circuit 1 but the long-range inhibitor in Circuit 2. The key difference is the regulatory logic that interprets BMP4 input and controls its production: BMP4 activates its own expression in Circuit 1, but induces a synthetic miRNA that represses the SNIPR activator ligand in circuit 2. Thus, signaling channels and intracellular logic can be swapped between modules, demonstrating that circuit components can be chosen for communication range and assigned network roles by wiring.

Model–experiment comparison provides one possible explanation for why our circuits, like other synthetic LALI patterning systems (*9*, *10*, *35*), generate patterns with a characteristic spatial scale but lack the long-range order associated with deterministic Turing patterns. In the model, transcriptional heterogeneity alone is sufficient to qualitatively reproduce irregular features of the experimental patterns (Fig. 3B and Fig. 5C). This indicates that irregularity need not reflect incomplete circuit optimization but may emerge from a minimal LALI circuit operating in a heterogeneous cellular context and suggests that the precise long-range order observed in natural patterning processes likely requires additional noise-suppressing or error-correcting mechanisms beyond the core LALI circuit. Conversely, although this heterogeneity reduced long-range order, it also broadened the pattern-forming regime in our model, consistent with previous work on bacterial synthetic Turing patterns (*10*).

The present circuits provide a proof of concept for patterning with a modular architecture that decouples communication from computation. However, several challenges for synthetic patterning circuits remain. First, in a closed cell culture system, the accumulation of secreted factors reduces spatial heterogeneity in signal distributions over time and saturates cellular responses. Addressing this problem in complex systems may require advances in cell-line engineering to enable active signal clearance, while microfluidic perfusion offers a more immediate route to continuous signal clearance in culture. On the other hand, in the organismal context, natural turnover mechanisms may make these issues less important and allow steady-state patterning. Second, the spatial information generated by these circuits must be made interpretable by downstream circuitry. Unlike the idealized steady-state patterns often considered in classical mathematical models, reaction–diffusion patterns in biological systems need not reach a final steady state or persist indefinitely. They need only establish interpretable spatial information within a finite time window, which can then be locked in by downstream stabilization circuits such as MultiFate (*36*) to produce durable compartment identities. Third, a broader repertoire of diffusible bio-orthogonal signals would facilitate engineering of more complex synthetic patterning systems.

More broadly, multicellular development represents an astonishingly complex, dynamic, and yet predictable biological program that critically depends on the ability to break symmetries and establish self-organized spatial patterns. This work complements synthetic and stem-cell-based approaches that pattern tissues through imposed morphogen gradients, geometry, adhesion, mechanics, or programmed cell assembly (*37–45*). By showing how autonomous LALI circuit logic can be built from defined mammalian signaling modules, these circuits should advance the rational programming of dynamic multicellular development.

## Methods

### Plasmid Construction

Plasmids were constructed following standard molecular biology protocols. Plasmid backbones were linearized using New England Biolabs (NEB) restriction enzymes. DNA inserts were generated by PCR or gBlock synthesis (IDT), and ligated into backbones using either standard Quick Ligation (NEB) or Gibson Assembly. Sequences were confirmed right before cell culture experiments using Plasmidsaurus whole-plasmid sequencing service.

### Cell Culture

The cell lines used in these experiments were AD-293 (Agilent Cat# 240085), a derivative of the standard HEK293 cell line selected for improved adherence and plaque formation, and U2OS (ATCC Cat# HTB-96), a widely used human osteosarcoma line characterized by robust adhesion, flat morphology, and suitability for imaging-based assays. All cell lines used in subsequent experiments were derived from one of these parental lines.

For routine maintenance, cells were cultured at 37 °C in a humidified incubator with 5% CO₂ in Dulbecco’s Modified Eagle Medium (DMEM) supplemented with 10% fetal bovine serum (FBS), 1× penicillin–streptomycin–L-glutamine, 1 mM sodium pyruvate, and 1× non-essential amino acids (NEAA). Cells were passaged every three days at a ratio of 1:10.

For maintenance of cells containing the BMP4 self-activation circuit, recombinant Noggin (250 ng/mL) was included in the culture medium to suppress spontaneous pathway activation and maintain the circuit in an “off” state.

For imaging experiments, the culture medium was replaced with FluoroBrite™ DMEM (Gibco) to reduce background fluorescence. Where indicated, 0.1 μM TMP and 5 μM Palbociclib were added to the medium to modulate fluorescent protein stability and cell cycle progression, respectively.

### Cell line construction

AD-293 cells were seeded in 24-well plates at 200,000 cells per well and cultured overnight prior to transfection. Transfection was performed using Lipofectamine LTX (Invitrogen #A12621) according to the manufacturer’s instructions. U2OS cells were seeded in 24-well plates at 50,000 cells per well and cultured overnight prior to transfection. Transfection was performed using Fugene HD (Promega #E2311) following the manufacturer’s protocol.

For stable cell line generation, cells were co-transfected with plasmids encoding the desired constructs together with a PiggyBac transposase expression vector. Following transfection, cells were subjected to antibiotic selection using one or more of the following: hygromycin (300 µg/mL), blasticidin (10 µg/mL), and puromycin (3 µg/mL), for at least two passages to enrich for stably integrated populations. To obtain homogeneous populations, monoclonal cell lines were established from the selected cells by limiting dilution or fluorescence-activated cell sorting (FACS).

### Flow Cytometry

Cells were seeded in a 96-well plate at a density of 30,000 per well and induced for 2 days before analysis. Cells were trypsinized with 0.05% trypsin for 5 minutes at room temperature (for AD-293) or 0.25% trypsin for 5 minutes at 37 °C. After digestion, cells were resuspended with HBSS (1x Hank’s Balanced Salt Solution) containing 2.5 mg/mL BSA (Bovine Serum Albumin) and 2 mM MgCl_2_ (Magnesium Chloride), filtered through a 40-μm cell strainer, and analyzed using a CytoFLEX S instrument (Beckman Coulter). Flow cytometry data were subsequently analyzed with EasyFlow (https://github.com/AntebiLab/easyflow).

### Cell Sorting Procedures

Cells were harvested post-trypsinization, resuspended in the sorting buffer containing HBSS supplemented with 2.5 mg/mL BSA, 2 mM MgCl_2_, and 1U/mL DNase I, and filtered through a 40-μm cell strainer before loading on the cell sorter (Sony MA900). Cells were sorted as mono-clones into 384-well plates or 96-well plates containing regular culture media.

### Reconstitution of Morphogen Gradients

Poly-D-lysine (0.1 mg/mL) was mixed with DPBS at 1:1 ratio. 1 mL of the mixture was loaded in each well of a 6-well plate and incubated at room temperature for 1 hour. The coating solution was then removed, and the plate was rinsed three times with 1.5 mL of ddH_2_O, dried, sealed, and stocked at 4 °C for up to two weeks before using. An iBiDi insert (#80209) was carefully positioned and gently pressed at the center of the well.

Senders were trypsinized, resuspended, and seeded at 80k per 100 µL media for each iBiDi insert. After 12 to 24 hours post-seeding, the iBiDi insert was carefully removed with tweezers and rinsed once with 1 mL media. 0.6 million receiver cells were then loaded in 2 mL media for each well. After receivers settled down, media was removed and the well was rinsed once with 1 mL imaging media. A mix of 1.5 mL imaging media containing small-molecule inducers and 0.5 mL Matrigel were loaded on cells. The plate was settled in the incubator for 15 min and transferred to the EVOS incubator (OSI Thermo) afterwards for time-lapse movies.

48 hrs of time-lapse movies were taken and the samples were scanned every 30 min. For each well, 10×5 field of views (FOVs) were scanned, with phase channel adjusting the autofocus, BFP channel locating the sender-receiver boundary, and YFP or mCherry signal indicating the signal gradients.

### Turing Patterning Experiment Setup

100 µL 0.1 mg/mL poly-D-lysine solution was added to a well of a 96-well plate for 1 hr at room temperature. Afterwards, the plate was thoroughly washed with 200 µL ddH_2_O three times to remove any residual poly-D-lysine, and air-dried at room temperature for 24 hrs. For the first circuit, cells were cultured in 150 µL media with 250 ng/mL Noggin for 24 hrs and rinsed with 300 µL regular media before seeding; for the second circuit, cells were cultured in a Shield1-free media sparsely for at least 24 hrs before seeding, both to prevent pre-patterning. Patterning monoclonals were then seeded at full confluence into the prepared plates in 500 µL patterning media. Cells were placed under an environmental chamber-equipped Olympus microscope for time-lapse movie capture.

Time-lapse movies were taken by an inverted Olympus IX81 fluorescence microscope with an ASI 2000YX automated stage, an iKon-M charge-coupled device camera (Andor), a 20× dry objective lens (0.7 NA), and an X-Cite XLED1 light source (Lumen Dynamics). Microscope and image acquisition were controlled by Metamorph software (Molecular Devices). Cells were kept in a customized environmental chamber with a humidified, 37 °C and 5% CO2 atmosphere. For each well, a square area at the well center was imaged. The region consists of 25 FOVs (5×5), and each FOV is 650 µm × 650 µm, with a 50-µm overlap between two adjacent FOVs. Movies were started 1 hour after induction and taken every 1 hour.

### Image preprocessing and pattern-spacing quantification

Pattern spacing was quantified from independent time-lapse movies after image preprocessing and radial spectral analysis, as described in Supplementary Methods. Briefly, each frame was corrected for slowly varying background intensity, robustly normalized, and analyzed by radial Fourier transform and radial autocorrelation. The primary pattern-spacing estimate reported in the main text is the centroid of the dominant radial FFT peak, with FFT max-bin, twice the first radial-autocorrelation minimum, and the first positive radial-autocorrelation peak retained as secondary estimates for method comparison. The statistical unit was an independent time-lapse movie. Sample sizes were circuit 1 patterning condition, n = 5 movies; circuit 1 self-activation-only control, n = 4 movies; and circuit 2 patterning condition, n = 4 movies. Unless otherwise indicated, values are reported as mean ± sample s.d.

## Acknowledgement

We thank Yunqing Wang, Yutian Li, Kaiwen Luo, Yitong Ma, Ronghui Zhu, Leah Santat, Pulin Li, Mark Budde, Joseph Markson, LiAng Yao, Zitong (Jerry) Wang, and other previous and current Elowitz lab members for sharing materials and scientific feedback; Justin Bois, Jordi Garcia-Ojavo, Pranav Bhamidipati, and Matt Thomson for inspiring discussions and technical support; Ethan Richman for providing critical feedback on the manuscript.

## Funding

This work was funded by the National Institutes of Health under award number R01HD075335, Allen Discovery under award number UWSC10142, and Sloan Foundation grant G-2024-22436.

## Author contributions

S.W., R.D., and M.B.E. conceived the project. S.W., R.D., and J.M.L performed all the experiments and analyzed the data. S.W., R.D., J.M.L., and M.B.E wrote the manuscript.

## Competing interests

The authors declare no competing interests for the work in this paper.

## Data, code, and materials availability

All data and custom code supporting the findings of this study are available through the CaltechDATA repository at https://doi.org/10.22002/x9259-tr458. Plasmids and cell lines are available from M.B.E. upon reasonable request.

## Notes

https://doi.org/10.22002/x9259-tr458

### Competing Interest Statement

The authors have declared no competing interest.

