## Supplementary Material for "Synthetic reaction-diffusion patterning in mammalian cells"

### Supplementary Materials

#### 1 Engineering and quantification of sender–receiver communication modules

The two synthetic LALI circuits are built from three intercellular communication modules: contact-dependent SNIPR–EGFRvIII signaling, diffusible BMP4 signaling, and diffusible Activin signaling. We first established sender–receiver formats for each module so that their spatial modes of communication and their corresponding reporter outputs could be compared in a common geometry (fig. S1).

In the quasi-one-dimensional sender–receiver assay, a removable insert initially confines the sender population. After sender cells attach, the insert is removed and receiver cells are plated adjacent to the sender region, producing a continuous monolayer with a defined sender–receiver interface (fig. S1A). Reporter fluorescence can then be measured as a function of distance perpendicular to this interface. This geometry was used for the signaling-spread measurements in fig. S2.

The genetic implementations of the three modules are summarized in fig. S1B. For the SNIPR–EGFRvIII module, sender cells constitutively display the EGFRvIII ligand, whereas receiver cells express SNIPR and a UAS reporter. Ligand engagement activates SNIPR and releases the Gal4–VP64 transcriptional activator, thereby inducing UAS-driven reporter expression. Because this module requires sender–receiver contact, it provides the local communication channel used in Circuit 2.

For the two diffusible modules, BMP4 or Activin production is induced specifically in sender cells. BMP4 sender cells use an ABA-responsive split Gal4-VP64 system to drive UAS-BMP4 expression. In receiver cells, BMP4 signaling is read out by phosphorylation of Smad1/5/8 and a BMP response element (BRE)-driven mCherry reporter. Activin sender cells use a doxycycline-responsive rtTA/TRE system to drive Activin expression. In receiver cells, Activin signaling is read out by phosphorylation of Smad2/3 and an Activin response element (ARE)-driven mCitrine reporter. Thus, fig. S1 defines the communication and reporting interfaces used throughout the study; their effective signaling spread is quantified separately in fig. S2.

#### 2 Estimation of effective signaling-spread coefficients from time-lapse movies

To compare the spatial propagation of BMP4- and Activin-dependent signaling, we quantified an effective signaling-spread coefficient from time-lapse reporter movies acquired using an EVOS imaging system (fig. S2). This coefficient describes the rate at which reporter-defined signaling activity expands within the receiver-cell layer.

For Activin, we used a corrected movie in which fields of view with unstable illumination were corrected using intensity references from adjacent frames, and a stage displacement between frames 58 and 59 was corrected by translational registration. For BMP4, we used an uncorrected movie and did not apply additional registration.

Before profile extraction, images were downsampled to 20% of their original width and height, corresponding to a fivefold reduction in each spatial dimension. Given the original pixel size of 1.725 $\mu$m/pixel, the effective pixel size after downsampling was:

$$\frac{1.725}{0.2} = 8.625\ \mu\text{m/pixel}. \quad (1)$$

For each movie and time point, a one-dimensional fluorescence profile was generated by averaging reporter fluorescence intensity along the $y$-axis (fig. S2, A and B). Analysis was restricted to a fixed distal receiver-only region located away from the sender–receiver boundary, where sender-cell contamination and local cell-density variation could otherwise bias the measured profiles. The analysis regions are indicated by orange boxes in the kymographs (fig. S2A). Within each condition, the same analysis start position was used across all five movies. The left edge of this analysis window was defined as $x = 0$ for the moment calculation.

Background fluorescence was estimated independently for each movie and time point and subtracted from the corresponding one-dimensional profile. Negative values after background subtraction were set to zero. The resulting profiles were smoothed along the $x$-axis using a Gaussian smoothing window. Representative fluorescence profiles at selected time points are shown in fig. S2B. Because the reporters used here have short half-lives, their fluorescence provides an approximate readout of recent pathway activity.

For each time point, we calculated the reporter-weighted second spatial moment:

$$M_2(t) = \frac{\sum_x I(x,t)x^2}{\sum_x I(x,t)}, \tag{2}$$

where $I(x,t)$ is the background-subtracted reporter fluorescence intensity at position $x$ and time $t$. Because this moment is calculated relative to the fixed start of the distal analysis window, it captures both broadening of the reporter profile and its progressive displacement toward more distal receiver positions.

The effective signaling-spread coefficient, $D_{\text{eff}}$, was defined from the approximately linear increase in $M_2(t)$ (*46, 47*):

$$M_2(t) = 2D_{\text{eff}}t + C, \tag{3}$$

where $C$ is a constant. Thus,

$$D_{\text{eff}} = \frac{1}{2}\frac{dM_2}{dt}. \tag{4}$$

For each ligand, the fitting interval was selected after reporter signal had become established in the distal receiver region and before the trajectory approached a plateau or was substantially affected by the finite imaging window. Activin entered this approximately linear regime earlier than BMP4, consistent with its faster signaling propagation. Frames 21–50 were used for Activin, whereas frames 51–80 were used for BMP4. Each frame interval corresponded to 1800 s. The $M_2(t)$ trajectories and corresponding linear fits are shown in fig. S2C.

This analysis produced one empirical estimate of $D_{\text{eff}}$ for each movie. Reported values represent the mean across five movies, with variability reported as the sample standard deviation (fig. S2D).

### 3 Specificity and separability of BMP4/BRE, Activin/ARE, and SNIPR/UAS signaling responses

The two circuit designs require multiple signaling modules to operate in the same cells. We therefore characterized their input–output relationships directly rather than assuming complete molecular orthogonality (fig. S3). Circuit 1 requires BMP4/BRE and Activin/ARE responses that are dominated by their intended ligands, whereas Circuit 2 requires a contact-dependent SNIPR/UAS response that remains compatible with BMP4/BRE signaling.

To examine the two TGF-$\beta$-family reporter modules, AD-293 cells carrying both BRE-driven mCherry and ARE-driven mCitrine were exposed to a matrix of recombinant BMP4 and Activin A concentrations (fig. S3A). BRE output increased primarily across the BMP4 axis, whereas ARE output increased primarily across the Activin axis. Neither reporter was completely insulated from the other ligand or from shared cellular context (*48*); however, over the concentration range examined, each reporter was dominated by its intended input. These measurements therefore establish functional input selectivity suitable for the BMP4/Activin circuit, rather than strict pathway orthogonality.

We next assessed whether SNIPR/UAS signaling and BMP4/BRE signaling could function in the same receiver-cell context (fig. S3B). UAS output was high only in the presence of EGFRvIII sender cells, consistent with the required juxtacrine input to SNIPR. Recombinant BMP4 did not induce UAS output in the absence of EGFRvIII senders. Conversely, BMP4 increased BRE output in both sender conditions, showing that

BMP4 responsiveness was retained in the SNIPR receiver-cell background. The magnitude of the two outputs varied with the combined signaling context, so these data should be interpreted as evidence of compatibility and the intended regulatory directions, not as evidence that the channels are fully independent.

Together, these measurements support the design logic used in the full circuits: BMP4 predominantly drives BRE output, Activin predominantly drives ARE output, and EGFRvIII sender contact drives SNIPR/UAS output. This level of input selectivity is sufficient to assign the required regulatory roles by circuit wiring while recognizing that the components share a common cellular environment.

### 4 Open-loop validation of LALI regulatory modules

Before combining the branches into closed spatial-feedback circuits, we tested whether the relevant regulatory relationships were functional in open-loop cell lines (fig. S4). These measurements test the direction of the engineered regulatory interactions in a uniform culture setting. They do not by themselves establish the relative signaling ranges that generate LALI behavior; those ranges are addressed by the sender–receiver assays and signaling-spread measurements in figs. S1 and S2.

For Circuit 1, the BRE cassette coexpresses BMP4 and mCherry, whereas the ARE cassette coexpresses Smad6 and mCitrine (fig. S4A). In the presence of TMP, which stabilizes the DHFR-tagged reporter, the line showed high mCherry output without added BMP4. Thus, exogenous BMP4 was not required to reveal activity of the self-activation module under these conditions. Addition of Activin A progressively reduced mCherry output over the displayed dose series. This result is consistent with the intended regulatory path in Circuit 1: Activin-responsive expression of Smad6 can suppress the BMP-responsive activation branch.

For Circuit 2, the UAS cassette coexpresses the EGFRvIII feedback ligand and mCherry, and the BRE cassette coexpresses mCitrine and the synthetic microRNA that targets the EGFRvIII transcript (fig. S4B). TMP stabilized the reporter, and Shield1 increased UAS-associated mCherry output, consistent with stabilization of the SNIPR receptor and activation of the juxtacrine positive-feedback branch. The displayed BMP-pathway perturbation series shifted mCherry output relative to the Shield1-stabilized condition, indicating that the SNIPR branch is responsive to modulation of the BMP-responsive regulatory module.

Collectively, the open-loop assays verify that the two architectures contain the regulatory branches needed for their subsequent closed-loop implementations: a BMP-responsive self-activation module that can be attenuated through the Activin–Smad6 arm in Circuit 1, and a Shield1-tunable SNIPR activation branch coupled to a BMP-responsive microRNA module in Circuit 2. The full-circuit patterning experiments test how these branches interact across space.

### 5 TMP-inducible reporter system

To track circuit dynamics during pattern formation, we designed a fluorescent reporter whose stability can be regulated post-translationally with trimethoprim (TMP) (fig. S5A). Each reporter was encoded on the same bicistronic transcript as its corresponding patterning component, with an internal ribosome entry site (IRES) enabling independent translation. This arrangement ensures that reporter production is driven by the same promoter as the associated circuit module, while avoiding direct fusion to the functional circuit protein.

To minimize fluorescence accumulated before the start of the experiment, the reporter was fused to H2B and a TMP-responsive DHFR destabilization domain. In the absence of TMP, the DHFR degron promotes rapid degradation of the reporter, preventing substantial fluorescent-protein accumulation even when basal promoter activity occurs during cell expansion or plating. Addition of TMP stabilizes the DHFR-tagged reporter, allowing newly synthesized fluorescent protein to accumulate to detectable levels. Thus, fluorescence primarily reflects reporter produced after TMP addition rather than residual protein synthesized before patterning was initiated.

We quantified the effect of TMP concentration on reporter stability using mCitrine (fig. S5B). Reporter fluorescence decayed approximately exponentially after TMP withdrawal, and the apparent half-life increased with TMP concentration. At 0.03, 0.1, 0.3, and 1.0 $\mu$M TMP, the measured mCitrine half-lives were approximately 8.5, $12.4 \pm 0.8$, 23.4, and 33.0 h, respectively. In the absence of TMP, fluorescence remained near background levels and a reporter half-life could not be reliably determined. TMP therefore provides a tunable reporter-memory window: lower TMP concentrations allow the reporter to respond more rapidly to recent changes in promoter activity, whereas higher concentrations increase signal accumulation and detection sensitivity.

Unless otherwise indicated, patterning experiments were performed in the presence of 0.1 $\mu$M TMP. Under this condition, the reporter half-life of approximately 12 h provided sufficient signal for long-term imaging while limiting excessive temporal averaging that would obscure dynamic changes during pattern formation. In the two patterning circuits, mCherry and mCitrine reported the activities of the short-range and long-range signaling branches, respectively, enabling simultaneous visualization of the spatial and temporal evolution of local activation and longer-range inhibition.

### 6 Clone construction and screening

We used a staged, clone-based construction strategy to access the expression configurations required for effective pattern formation (fig. S6A). Circuit modules were introduced sequentially by piggyBac transposition. Because piggyBac integration is stochastic, individual clones differ both in transgene copy number and in the local chromatin environment of their genomic integration sites, generating distinct effective expression levels across circuit modules. Rather than treating this clone-to-clone variation solely as technical heterogeneity in stable cell-line construction, we used it to sample the parameter space defined by the relative expression of the activation and inhibition modules.

This strategy differs from conventional stable cell-line construction, in which the objective is typically to obtain uniform expression across clones. Here, pattern formation is expected to occur only within a restricted region of parameter space; therefore, not every monoclonal line is expected to pattern. PiggyBac-based random integration provides an efficient means to generate and test diverse circuit-expression configurations without independently tuning each circuit component. All cell lines used for subsequent experiments and analyses were monoclonal lines derived by single-cell sorting (fig. S6A).

We first introduced the BMP-responsive self-activation module, followed by antibiotic selection, single-cell sorting, and clonal expansion (fig. S6B, left). The purpose of this first screen was not to identify pattern-forming clones directly, but rather to select a parental clone with robust and reproducible BRE self-activation. Multiple monoclonal lines showed detectable self-activation, from which we selected a clone with strong and stable BRE output as the parental line for the second construction step.

We next introduced a second plasmid encoding the Activin-responsive inhibitory module into this parental clone, followed by a second round of antibiotic selection and single-cell sorting (fig. S6B, right). The resulting monoclonal lines were screened for dual reporter expression under TMP-containing reporter-stabilizing conditions, but without compounds used to perturb or tune circuit parameters. We selected clones with detectable signals from both the BRE reporter, mCherry, and the ARE reporter, mCitrine, thereby identifying candidate lines in which both the activation and inhibitory branches were integrated and expressed. Clones with strongly imbalanced reporter expression, for example, high mCherry with low mCitrine or the converse, were deprioritized because such imbalance was unlikely to support the intended coupled-feedback architecture.

Among the monoclonal lines obtained in the second screen, 13 clones showed relatively balanced and robust mCherry and mCitrine signals and were subsequently evaluated in standardized patterning assays (fig. S6B). The primary experiments and quantitative analyses in this study were performed using clone 22J.

### 7 Patterning in additional clones

To assess how patterning behavior varied across independently derived full-circuit monoclonal lines, we performed time-lapse imaging of four additional clones under the same standardized patterning conditions used for the primary characterization (fig. S14). All clones were derived by single-cell sorting after sequential piggyBac integration of the self-activation and inhibitory modules. Because piggyBac integration produces clone-specific transgene copy numbers and genomic integration contexts, individual monoclonal lines are expected to occupy different effective expression regimes within the circuit parameter space.

The additional clones displayed a range of dynamic behaviors. Clones 9H and 22D developed localized mCitrine-positive domains that became more pronounced over time, with mCherry signal emerging in subsets of the field at later stages. Clone 12K showed progressively heterogeneous dual-reporter activity, but with less clearly resolved spatial domains. In contrast, clone 15D showed a broad increase in mCitrine signal with comparatively limited spatial segregation. Thus, independently derived monoclonal lines did not converge on a single common morphology; instead, they sampled a continuum of circuit behaviors ranging from relatively homogeneous activation to localized, domain-like patterning.

These results are consistent with the intended clone-selection strategy. In this system, piggyBac-mediated integration was used to generate a diverse set of relative expression configurations rather than uniformly equivalent cell lines. Accordingly, independently derived full-circuit clones were expected to sample a continuum of circuit behaviors, and not every monoclonal line was expected to support robust pattern formation. Clone 22J was selected for detailed analysis because it reproducibly produced well-resolved patterns with sufficient contrast and temporal stability for quantitative characterization, whereas the additional clones shown here illustrate the broader diversity of outcomes generated by the same circuit architecture.

### 8 Active-domain coarsening analysis

Following established connected-domain and coarsening analysis strategies (*49, 50*), we tested whether the patterning circuit changed the spatial organization of active signaling domains over time, rather than only changing the total fluorescence intensity. If local activation domains coarsen during pattern formation, small domains should become less frequent and the largest connected domains should become larger. We therefore converted fluorescence images into binary active-domain maps and quantified connected active regions at multiple time points.

For circuit 1, we analyzed mCherry movies at 15, 30, 45, and 60 h. Movies with 31 frames were sampled at frames 6, 11, 16, and 21, and the 42-frame movie was sampled at frames 11, 21, 31, and 41. Each image was first smoothed with a Gaussian kernel of $\sigma = 8.8$ $\mu$m, corresponding to 1 pixel because the image pixel size was 8.8 $\mu$m per pixel. The smoothed image was then thresholded at the 85th percentile, so that the brightest 15% of pixels were called active. Connected foreground regions were identified with 4-connectivity, and domains smaller than 1,000 $\mu m^2$ were excluded from the primary domain-size distribution. For each movie and time point, we recorded the area of every retained connected domain and summarized both the number of active domains and the distribution of large-domain areas. These measurements produced the circuit 1 active-domain quantification shown in Fig. 2D.

Because the exact domain calls depend on image smoothing and thresholding, we treated the segmentation parameters as an analysis caveat and tested whether the qualitative result was robust to reasonable parameter choices. We screened Gaussian smoothing sigmas of 4.4, 8.8, and 13.2 $\mu$m and intensity thresholds at the 80th, 85th, and 90th percentiles, corresponding to the brightest 20%, 15%, and 10% of pixels. fig. S7B summarizes this parameter screen as fold change relative to the first time point. In each heatmap, the first row is the circuit 1 patterning condition and the second row is the self-activation-only control; the four columns are the four time points. Values were normalized to the first time point within each row, so red indicates an increase over time and blue indicates a decrease. Across the screen, circuit 1 showed a decrease in the number of small active domains and an increase in the size of the largest domains, whereas the self-activation-only control showed the opposite or much weaker trend.

For the small-domain metric, we counted active domains with areas between 1,000 and 10,000 $\mu m^2$. For the large-domain metric, we simplified the tail of the domain-area distribution by taking the median area of the largest domains rather than relying on a single maximum value; in the screen we evaluated the median of the top 10, top 20, and top 30 domains. These robustness summaries are shown in fig. S7, C and D. The same active-domain workflow was then applied to circuit 2 movies, using the same frame-selection, smoothing, thresholding, connected-component, and area-summary procedure, producing the circuit 2 analysis in fig. S11D.

### 9 Domain-centered superposition analysis

To compare the local spatial relationship between the red and green channels, we performed domain-centered LALI profiling by superimposing image neighborhoods centered on red-channel active domains. We first chose a Laplacian-of-Gaussian (LoG) filter scale matched to the pattern length scale estimated from the spatial correlation function $C(r)$ (*51*). Operationally, we scanned candidate LoG sigmas on the 60 h mCitrine images and selected a single group-level filter scale from the scale-normalized LoG peak response. For circuit 1, the candidate sigmas were 3, 4, 5, 6, 8, 10, 12, 15, 18, and 20 pixels, and $\sigma = 10$ pixels was selected. Images were clipped between the 1st and 99th percentiles before filtering to reduce the influence of rare bright outliers.

Using this selected filter scale, we applied the scale-normalized LoG filter to the corresponding 60 h mCherry images and detected local maxima as red-domain centers. Candidate centers were required to exceed mean + 1 SD of the LoG response, to be separated by at least $2\sigma$ during peak detection, and to lie far enough from the image border to extract a complete neighborhood. After candidate detection, centers were greedily pruned by LoG response so that retained centers were at least two neighborhood radii apart. For circuit 1, the selected sigma gave a neighborhood radius of 30 pixels, corresponding to 264 $\mu$m, and a minimum center-to-center distance of 60 pixels. We then extracted matched mCherry and mCitrine patches around each retained center, averaged the patches across centers and movies, and computed radial-average fluorescence profiles. As a spatial control, we shuffled the row and column coordinates of the detected centers while preserving their marginal distributions, then repeated the same patch averaging and radial profiling.

The resulting superimposed average in Fig. 2E shows that the red signal is concentrated near the center of the detected domains, whereas the green signal extends over a broader radial range. This red-narrow/green-wide organization is consistent with a local-activation, long-range-inhibition (LALI) spatial relationship.

We applied the same mCherry-centered superimposed-average workflow to circuit 2 for fig. S11E. For circuit 2, the candidate LoG sigmas were 7, 8, 9, and 10 pixels, and $\sigma = 7$ pixels was selected, giving a neighborhood radius of 21 pixels, corresponding to 185 $\mu$m, and a minimum center-to-center distance of 42 pixels. The circuit 2 superimposed averages and radial profiles again showed a narrow red peak with a broader green profile, consistent with the same LALI organization observed for circuit 1.

### 10 Pattern spacing quantification

Pattern-spacing measurements were designed to capture the dominant spatial repeat scale of fluorescence patterns while reducing sensitivity to non-biological imaging artifacts. The same preprocessing and radial autocorrelation workflow was used for the main-figure autocorrelation analyses in Fig. 2F and Fig. 5B and for the replicate-level spacing audits in fig. S8 and fig. S11. Fig. 2F and Fig. 5B show the time-resolved radial-averaged autocorrelation profiles, $C(r)$, used to visualize the emergence of a characteristic length scale during pattern formation. fig. S8 and fig. S11 show the per-movie 60 h measurements used to compare spacing estimators for circuit 1 and circuit 2, respectively.

All analyses were performed on independent time-lapse movies, with each movie treated as one statistical unit. The sample sizes used for spacing summary statistics were circuit 1 patterning condition, $n = 5$ movies, corresponding to the five columns in fig. S8, A and B; circuit 1 self-activation-only control, $n = 4$ movies; and circuit 2 patterning condition, $n = 4$ movies, corresponding to the four columns in fig. S11, A and B. Values are reported as mean $\pm$ sample standard deviation unless otherwise indicated.

For each analyzed frame, the raw fluorescence image was converted to double precision and non-finite pixels, if any, were temporarily filled with the median finite-pixel value for filtering. Slow background structure was estimated by large-scale Gaussian blurring the image and subtracting this blurred image from the raw image. This background-subtraction step was used to correct uneven illumination, vignetting, and other low-spatial-frequency intensity offsets that would otherwise dominate the zero-frequency and low-frequency components of the FFT (*52*). The background-subtracted image was then lightly smoothed with a Gaussian filter with $\sigma = 8.8$ $\mu$m to suppress pixel-scale noise without removing the pattern-scale signal. For normalization across movies and time points, the corrected image was centered by its median finite-pixel intensity and divided by a robust scale estimate, $1.4826\times$ the median absolute deviation; if this robust scale was numerically degenerate, the standard deviation or 1 was used as a fallback. This preprocessing was applied before generating the radial autocorrelation curves in Fig. 2F and Fig. 5B and before the FFT and $C(r)$ analyses summarized in fig. S8 and fig. S11.

Radial autocorrelation was computed from the preprocessed image after subtracting the mean finite-pixel intensity. The two-dimensional autocorrelation was evaluated using FFT-based convolution with an accompanying binary mask autocorrelation to normalize for the number of overlapping pixels at each displacement. The autocorrelation map was normalized to its zero-lag value, and positions with less than 5% of the maximum overlap support were excluded. The radial autocorrelation profile, $C(r)$, was then obtained by averaging autocorrelation values in annular bins as a function of radial distance $r$. From $C(r)$, two spacing-related estimates were retained: twice the radius of the first minimum after the first zero crossing, and the radius of the first positive side peak after that minimum. The 2× first-minimum estimate reports the approximate separation between alternating high and low phases, whereas the first positive side peak directly estimates the repeat distance between positively correlated pattern features.

In the main figures, this $C(r)$ workflow is displayed as the radially averaged autocorrelation plots in Fig. 2F and Fig. 5B, where curves over 0–60 h show how the negative trough and subsequent positive side peak develop as the pattern matures. In the supplement, the same $C(r)$ profiles are shown for each independent movie in fig. S8B for circuit 1 and fig. S11B for circuit 2. The green dashed line marks the first negative minimum used for the 2× minimum estimate, and the brown dashed line marks the first positive side peak used for the direct peak-to-peak spacing estimate.

Radial FFT analysis was performed on the same preprocessed images. Finite pixels were mean-subtracted, non-finite regions were set to zero, and a two-dimensional cosine window was applied before FFT calculation to reduce edge discontinuities. The squared magnitude of the shifted two-dimensional FFT was radially averaged in frequency space using bins set by the fundamental spatial-frequency resolution of the image (*53, 54*). The resulting radial power spectrum was smoothed with a five-bin moving median. Candidate pattern spacings were searched over 40–1500 $\mu$m. The replicate-level 60 h radial FFT spectra are shown in fig. S8A for circuit 1 and fig. S11A for circuit 2. In these panels, the purple solid line marks the FFT max-bin estimate, the red solid line marks the FFT centroid estimate, and the shaded peak region indicates the dominant frequency lobe used for centroid calculation.

Four pattern-spacing estimates were calculated for auditability. First, the FFT max-bin estimate was defined as the pattern spacing corresponding to the frequency bin with maximum baseline-subtracted radial power within the search band. Second, the FFT centroid estimate was calculated from the dominant peak lobe: the lobe was bounded by the half-height crossings around the max bin, radial power was baseline-subtracted, and the centroid frequency was computed as the power-weighted mean frequency over positive lobe points; pattern spacing was then computed as the inverse of frequency. Third, the $C(r)$ 2× minimum estimate was defined as twice the radius of the first autocorrelation minimum after the first zero crossing. Fourth, the $C(r)$ first-peak estimate was defined as the radius of the first positive autocorrelation side peak after the first minimum.

For circuit 1, the four estimates are summarized in fig. S8C: FFT max-bin, $582.0 \pm 127.4$ $\mu$m; FFT centroid, $546.8 \pm 54.9$ $\mu$m; $C(r)$ 2× minimum, $475.2 \pm 21.6$ $\mu$m; and $C(r)$ first peak, $512.2 \pm 73.2$ $\mu$m. For circuit 2, the corresponding estimates are summarized in fig. S11C: FFT max-bin, $349.7 \pm 52.2$ $\mu$m; FFT centroid, $319.3 \pm 15.3$ $\mu$m; $C(r)$ 2× minimum, $250.8 \pm 26.4$ $\mu$m; and $C(r)$ first peak, $369.6 \pm 29.6$ $\mu$m. The main text reports the FFT centroid as the primary spacing estimate because it uses the full dominant FFT peak lobe rather than a single discrete bin, making it less sensitive to FFT bin placement while avoiding assumptions from parametric peak fitting. The remaining estimates were used as robustness checks that connect the main-figure $C(r)$ readouts in Fig. 2F and Fig. 5B to the full frequency-domain and real-space spacing audit in fig. S8 and fig. S11.

### 11 Lattice-based ODE models with cell-to-cell promoter heterogeneity

#### 11.1 Model framework

We simulated both synthetic LALI circuits using ordinary differential equations on a fixed two-dimensional square lattice, with each lattice node representing one cell. This formulation preserves cell-level differences in promoter activity while accommodating both diffusible extracellular signals and contact-dependent intercellular signaling. Model 1 represents the BMP4/Activin/Smad6 circuit, in which slowly diffusing BMP4 provides local activation and more rapidly diffusing Activin induces the intracellular BMP inhibitor Smad6. Model 2 represents the SNIPR/EGFRvIII/BMP4/miRNA circuit, in which contact-dependent SNIPR–EGFRvIII signaling provides short-range activation, whereas diffusible BMP4 activates a BRE-driven miRNA that represses EGFRvIII expression. Thus, BMP4 contributes to the activating arm in Model 1 but to the inhibitory arm in Model 2.

For each diffusible species $c$ at node $i$, diffusion was implemented using a discrete Laplacian over the four edge-adjacent neighbors (*55, 56*):

$$\frac{dc_i}{dt} = F(c_i, \ldots) + D_c \sum_{j \in N(i)} (c_j - c_i), \tag{5}$$

where $N(i)$ denotes the four edge-adjacent lattice nodes. Regulatory interactions were represented using Hill activation and repression functions:

$$f(x; K, h, \kappa) = \frac{(x/K)^h + \kappa}{(x/K)^h + \kappa + 1}, \tag{6}$$

$$g(x; K, h) = \frac{1}{(x/K)^h + 1}. \tag{7}$$

#### 11.2 BMP4/Activin/Smad6 circuit equations

The BMP4/Activin/Smad6 circuit was modeled as:

$$\frac{dB_i}{dt} = \beta_B f(B_i; K_B, H_B, \kappa_B) g(S_i; K_S, H_S) - \gamma_B B_i + D_B \sum_{j \in N(i)} (B_j - B_i), \tag{8}$$

$$\frac{dA_i}{dt} = \beta_A f(B_i; K_B, H_B, \kappa_B) g(S_i; K_S, H_S) - \gamma_A A_i + D_A \sum_{j \in N(i)} (A_j - A_i), \tag{9}$$

$$\frac{dS_i}{dt} = \beta_S f(A_i; K_A, H_A, \kappa_A) - \gamma_S S_i. \tag{10}$$

Here $B_i$, $A_i$, and $S_i$ denote BMP4, Activin, and Smad6 levels at lattice node $i$, respectively. BMP4 activates both BMP4 and Activin production through the BMP-responsive module. Activin induces Smad6 through the Activin-responsive module, and Smad6 inhibits BMP-responsive production. Thus, the model implements local activation and long-range inhibition through the slower diffusion of BMP4 and faster diffusion of Activin.

#### 11.3 SNIPR/EGFRvIII/BMP4/miRNA circuit equations

In Model 2, $E_i$, $B_i$, and $R_i$ denote EGFRvIII, BMP4, and the miRNA repressor at lattice node $i$, respectively. EGFRvIII is membrane tethered and therefore does not diffuse. Instead, EGFRvIII on neighboring cells activates SNIPR and the downstream UAS promoter. To represent this contact-dependent input, we calculated the mean EGFRvIII abundance among the four edge-adjacent neighbors and among all eight edge- and diagonally adjacent neighbors:

$$\overline{E}_i^{(4)} = \frac{1}{4} \sum_{j \in N_4(i)} E_j, \tag{11}$$

$$\overline{E}_i^{(8)} = \frac{1}{8} \sum_{j \in N_8(i)} E_j, \tag{12}$$

and combined these two neighborhood averages as

$$\overline{E}_i = (1 - \alpha)\overline{E}_i^{(4)} + \alpha \overline{E}_i^{(8)}. \tag{13}$$

Here $N_4(i)$ contains the four edge-adjacent nodes, $N_8(i)$ contains the four edge-adjacent and four diagonally adjacent nodes, and $\alpha = 0.5$ in the reference Model 2 simulations. This interpolation gives greater total weight to edge-adjacent contacts while retaining contributions from diagonal contacts on the square lattice.

The SNIPR/EGFRvIII/BMP4/miRNA circuit was modeled as:

$$\frac{dE_i}{dt} = \beta_E f(\overline{E}_i; K_E, h_E, \kappa_E) g(R_i; K_R, h_R) - \gamma_E E_i, \tag{14}$$

$$\frac{dB_i}{dt} = \beta_B f(\overline{E}_i; K_E, h_E, \kappa_E) - \gamma_B B_i + D_B \sum_{j \in N(i)} (B_j - B_i), \tag{15}$$

$$\frac{dR_i}{dt} = \beta_R f(B_i; K_B, h_B, \kappa_B) - \gamma_R R_i. \tag{16}$$

Activated SNIPR drives the UAS-dependent production of both EGFRvIII and BMP4. Neighboring EGFRvIII therefore reinforces contact-dependent UAS activation, whereas BMP4 diffuses between cells and activates BRE-dependent production of the miRNA repressor. The miRNA inhibits EGFRvIII production cell autonomously. BMP4 is the only diffusible species in Model 2; EGFRvIII acts through cell–cell contact and the miRNA repressor remains intracellular.

#### 11.4 Cell-to-cell promoter heterogeneity model

Heterogeneity among cells in promoter-driven expression was represented by a time-independent multiplicative factor on each production term. For a production term $\beta f$, we used:

$$\beta f \to \beta f \cdot \eta_i, \tag{17}$$

where $\eta_i$ is the factor assigned to the cell represented by lattice node $i$. The cell-to-cell production multiplier was modeled as a mean-corrected lognormal variable:

$$\eta_i = \exp\left(\sigma z_i - \frac{\sigma^2}{2}\right), \qquad z_i \overset{\text{iid}}{\sim} \mathcal{N}(0, 1). \tag{18}$$

This form ensures that:

$$E[\eta_i] = 1, \tag{19}$$

so that cell-to-cell heterogeneity does not change the mean production rate. In Model 1, a single cell-specific multiplier $\eta_i$ was applied to all three production rates, $\beta_B$, $\beta_A$, and $\beta_S$. Thus, $\sigma$ is the sole heterogeneity-amplitude parameter in Model 1. In Model 2, EGFRvIII and BMP4 were assigned the same multiplier because both are coexpressed from the UAS-driven module, whereas the miRNA repressor is expressed from the BRE-driven module and was assigned its own module-level multiplier.

#### 11.5 Parameter values and sources

The source or basis of each parameter is listed in the tables below. Figure citations identify values measured or derived from experiments in this study. Model-calibrated parameters were selected to define a reference pattern-forming regime; BioNumbers (*57*) identifiers indicate broad order-of-magnitude range checks rather than direct measurements of the corresponding model parameters. Assumed parameters and fixed reference scales were specified from circuit structure or model normalization.

#### 11.6 Parameter rationale

The lattice spacing was set to 20 $\mu$m, approximately one mammalian cell diameter and close to the cell spacing in the confluent monolayer experiments. This choice defines the spatial discretization of the lattice model rather than an additional fitted parameter.

The diffusion coefficients for BMP4 and Activin were measured from the one-dimensional signaling assay. The measured values were:

$$D_B = 1.65 \pm 0.17\ \mu\text{m}^2/\text{s}, \tag{20}$$

$$D_A = 10.03 \pm 0.77\ \mu\text{m}^2/\text{s}. \tag{21}$$

Table 1: Parameters used in the BMP4/Activin/Smad6 lattice-ODE model

| Parameter name | Symbol | Physical value | Nondim. value | Source / basis |
|---|---|---|---|---|
| Lattice spacing | $\Delta x$ | 20.0 $\mu$m | 1 | Model discretization; BNID 100432 |
| BMP4 diffusion coefficient | $D_B$ | $1.65 \pm 0.17\ \mu\mathrm{m}^2\,\mathrm{s}^{-1}$ | 14.88 | fig. S2D |
| Activin diffusion coefficient | $D_A$ | $10.03 \pm 0.77\ \mu\mathrm{m}^2\,\mathrm{s}^{-1}$ | 90.67 | fig. S2D |
| BMP4 production rate | $\beta_B$ | 26.4 $\mathrm{nM\,h^{-1}}$ | 26.51 | Model-calibrated; BNID 103954 range check |
| Activin production rate | $\beta_A$ | 19.8 $\mathrm{nM\,h^{-1}}$ | 19.88 | Model-calibrated; BNID 103954 range check |
| Smad6 production rate | $\beta_S$ | 30.0 $\mathrm{nM\,h^{-1}}$ | 30.12 | Model-calibrated |
| BMP4 removal rate | $\gamma_B$ | 0.996 $\mathrm{h^{-1}}$ | 1 | Model-calibrated; BNID 112252 range check |
| Activin removal rate | $\gamma_A$ | 0.498 $\mathrm{h^{-1}}$ | 0.50 | Model-calibrated; BNID 112252 range check |
| Smad6 removal rate | $\gamma_S$ | 9.90 $\mathrm{h^{-1}}$ | 9.94 | Model-calibrated; BNID 103760 range check |
| BMP4 activation threshold | $K_B$ | 1 nM | 1 | fig. S3A; $EC_{50}$-derived |
| Activin activation threshold | $K_A$ | 1 nM | 1 | fig. S3A; $EC_{50}$-derived |
| Smad6 repression threshold | $K_S$ | 1 nM | 1 | Fixed reference scale |
| BMP4 Hill coefficient | $H_B$ | 4 | 4 | Assumed |
| Activin Hill coefficient | $H_A$ | 2 | 2 | Assumed |
| Smad6 Hill coefficient | $H_S$ | 5 | 5 | Assumed |
| BMP4-dependent basal activation | $\kappa_B$ | 0.02 | 0.02 | Assumed |
| Activin-dependent basal activation | $\kappa_A$ | 0.02 | 0.02 | Assumed |
| Cell-to-cell production heterogeneity | $\sigma$ | 0.3 | 0.3 | Literature-informed reference magnitude (*26, 27*) |

Table 2: Parameters used in the SNIPR/EGFRvIII/BMP4/miRNA lattice-ODE model

| Parameter name | Symbol | Physical value | Nondim. value | Source / basis |
|---|---|---|---|---|
| Lattice spacing | $\Delta x$ | 20.0 $\mu$m | 1 | Model discretization; BNID 100432 |
| Edge/eight-neighbor mixing coefficient | $\alpha$ | 0.50 | 0.50 | Assumed |
| BMP4 diffusion coefficient | $D_B$ | $1.65 \pm 0.17$ $\mu\text{m}^2\,\text{s}^{-1}$ | 14.85 | fig. S2D |
| EGFRvIII production rate | $\beta_E$ | 12.0 $\text{nM}\,\text{h}^{-1}$ | 12.05 | Model-calibrated; BNIDs 104829/112720 range check |
| BMP4 production rate | $\beta_B$ | 12.0 $\text{nM}\,\text{h}^{-1}$ | 12.05 | Model-calibrated; BNID 103954 range check |
| miRNA repressor production rate | $\beta_R$ | 9.0 $\text{nM}\,\text{h}^{-1}$ | 9.04 | Model-calibrated |
| EGFRvIII removal rate | $\gamma_E$ | 0.222 $\text{h}^{-1}$ | 0.223 | Model-calibrated; BNID 112252 range check |
| BMP4 removal rate | $\gamma_B$ | 0.996 $\text{h}^{-1}$ | 1 | Model-calibrated; BNID 112252 range check |
| miRNA repressor removal rate | $\gamma_R$ | 2.748 $\text{h}^{-1}$ | 2.759 | Model-calibrated |
| SNIPR activation threshold | $K_E$ | 1 nM | 1 | Fixed reference scale |
| BMP4 activation threshold | $K_B$ | 1 nM | 1 | fig. S3A; $EC_{50}$-derived |
| miRNA repression threshold | $K_R$ | 1 nM | 1 | Fixed reference scale |
| SNIPR/UAS Hill coefficient | $h_E$ | 4 | 4 | Assumed |
| BMP4/BRE Hill coefficient | $h_B$ | 2 | 2 | Assumed |
| miRNA repression Hill coefficient | $h_R$ | 3 | 3 | Assumed |
| SNIPR-dependent basal activation | $\kappa_E$ | 0.02 | 0.02 | Assumed |
| BMP4-dependent basal activation | $\kappa_B$ | 0.02 | 0.02 | Assumed |
| Cell-to-cell production heterogeneity | $\sigma$ | 0.3 | 0.3 | Literature-informed reference magnitude (*26, 27*) |

These measurements indicate that Activin diffuses approximately six-fold faster than BMP4:

$$D_A/D_B \approx 6.1. \tag{22}$$

This measured transport asymmetry is the main experimentally constrained physical feature that enables Activin to act as the longer-range arm of the LALI circuit.

For Model 1, the single log-scale heterogeneity parameter was set to $\sigma = 0.300$ as a literature-informed reference magnitude rather than a value fitted to the reporter distributions in this study. Under the mean-corrected lognormal parameterization, $\sigma = 0.300$ corresponds to a linear-scale coefficient of variation of $\sqrt{\exp(\sigma^2) - 1} \approx 0.31$. Cell-to-cell protein-expression variability of this order has been reported in mammalian cells, including coefficients of variation of approximately 0.25–0.30 in synthetic expression constructs and standard deviations of approximately 15–30% of the mean for endogenous proteins (*26, 27*). These measurements motivate a representative moderate heterogeneity magnitude but do not directly validate the particular lognormal field or the value of $\sigma$ used here. The same cell-specific multiplier $\eta_i$ was applied to the BMP4, Activin, and Smad6 production rates. For the reference Model 2 simulations, the UAS-driven EGFRvIII/BMP4 module and the BRE-driven miRNA module each used a heterogeneity magnitude of 0.300.

Hill coefficients were treated as mechanistic assumptions rather than fitted quantities. The BMP4 self-activation Hill coefficient $H_B = 4$ represents cooperative BMP/SMAD signaling and transcriptional amplification. The Activin activation Hill coefficient $H_A = 2$ represents canonical Activin/SMAD-dependent activation. The Smad6 repression Hill coefficient $H_S = 5$ represents an ultrasensitive inhibitory step, consistent with Smad6-mediated suppression of BMP pathway output. The basal activation parameters $\kappa_B = \kappa_A = 0.02$ allow weak basal production in the absence of full pathway activation.

For Model 2, the SNIPR/UAS Hill coefficient $h_E = 4$ represents the effective cooperativity of contact-dependent receptor activation, transcription-factor release, and downstream transcriptional amplification. The BMP4/BRE Hill coefficient $h_B = 2$ represents BMP/SMAD-dependent activation of the BRE module, and $h_R = 3$ represents the nonlinear repression of EGFRvIII expression by the miRNA module. The basal activation parameters $\kappa_E = \kappa_B = 0.02$ allow weak expression in the absence of full pathway activation. Both models were integrated in nondimensional form, with concentration scaled by the corresponding $K_B = 1$ nM, time by $\gamma_B^{-1}$, and length by the lattice spacing $\Delta x = 20$ $\mu$m.

The production rates, removal rates, and regulatory thresholds were selected within biologically plausible ranges to define a reference model that lies in a pattern-forming regime. These parameters were not independently fit to each experimental perturbation condition. For perturbation simulations, only the signaling parameter corresponding to the perturbation was changed, while the rest of the reference parameter set was kept fixed. Therefore, perturbation trends should be interpreted as predictions from the calibrated reference model, rather than post-hoc fits to each perturbation condition.

##### 11.7 Simulation procedure

For each simulation, the corresponding homogeneous zero-diffusion ODE system was first solved to obtain a uniform steady state. This steady state was assigned to all lattice nodes. At $t = 0$, a small independent Gaussian perturbation was applied to each species at each node to break symmetry; the final scripts used a perturbation with a standard deviation equal to 2% of the local steady-state value. Cell-specific promoter multipliers were then assigned according to the heterogeneity model above, and the full lattice system was integrated using MATLAB's `ode15s` solver until the stopping criterion was reached. Reflective boundary conditions were used for diffusion and for the neighborhood averages that define contact-dependent SNIPR input. Simulations used a lattice spacing of 20 $\mu$m. When pattern statistics were sensitive to edge effects, simulations were performed on a larger lattice than the reported analysis region and quantification was restricted to the interior.

##### 11.8 Pattern quantification

Final simulated patterns were analyzed using radial autocorrelation and two-dimensional Fourier analysis. The radial autocorrelation function $C(r)$ was used to quantify short-range spatial organization and long-range order. The first off-center peak position of $C(r)$ was used as an estimate of the local pattern spacing, and the decay of the autocorrelation envelope was used to assess the loss of long-range spatial order.

In parallel, the radially averaged two-dimensional Fourier spectrum was used to identify the dominant nonzero spatial frequency $q^*$. The corresponding characteristic pattern spacing was calculated as:

$$\lambda = \frac{2\pi}{q^*}. \tag{23}$$

##### 11.9 Model limitations

The models do not include cell division, cell migration, changes in cell shape, or density-dependent mechanical effects. The square lattice approximates a confluent monolayer but does not capture the exact geometry of cell packing. In Model 2, cell–cell contact is represented by weighted neighborhood averages rather than by explicit receptor–ligand binding at measured cell boundaries. Intrinsic molecular fluctuations were not simulated explicitly; instead, heterogeneity among cells in promoter-driven expression was represented by fixed multiplicative factors.

The models should therefore be interpreted as reference models that test whether the implemented LALI circuit topologies, measured transport properties, and representative cell-to-cell production heterogeneity are sufficient to reproduce the main qualitative features of the observed patterns, rather than as uniquely identifiable fits of all biochemical parameters.

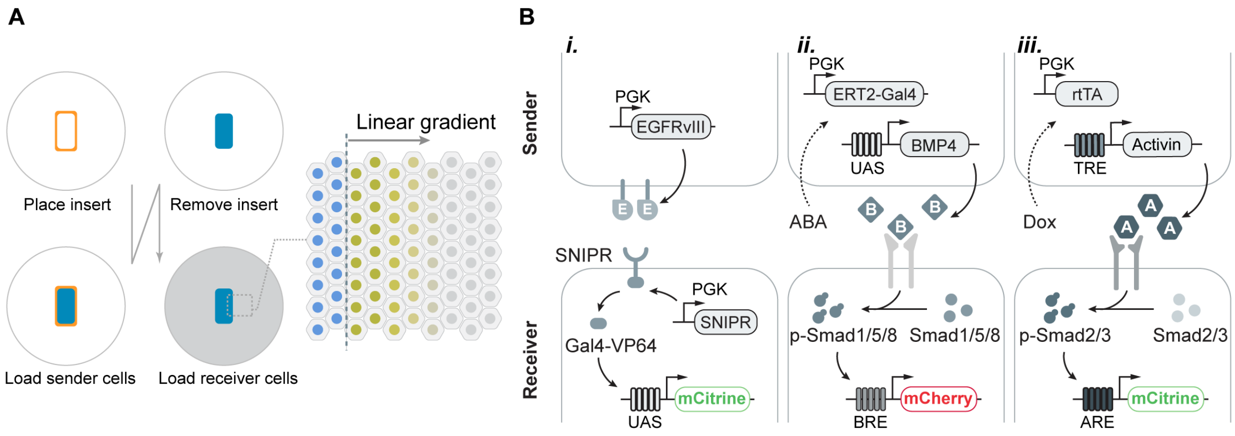


**Fig. S1. Sender–receiver assay and circuit architectures used to compare communication modules. (A)** Schematic of the quasi-one-dimensional sender–receiver assay. Sender cells were first confined within a removable insert (orange outline). After insert removal, receiver cells were plated to form a sharp sender–receiver interface, enabling signaling activity to be quantified as a function of distance from the sender region. **(B)** Sender and receiver constructs for the three communication channels. *i*, Juxtacrine SNIPR–EGFRvIII signaling: sender cells constitutively express membrane-tethered EGFRvIII, and direct sender–receiver contact activates SNIPR-dependent Gal4-VP64 release and a UAS-driven mCitrine reporter in receiver cells. *ii*, BMP4 signaling: ABA induces split Gal4-VP64 activity in sender cells to drive UAS-BMP4 expression; BMP4 activates the Smad1/5/8-responsive BRE-mCherry reporter in receiver cells. *iii*, Activin signaling: doxycycline induces rtTA/TRE-driven Activin expression in sender cells; Activin activates the Smad2/3-responsive ARE-mCitrine reporter in receiver cells.

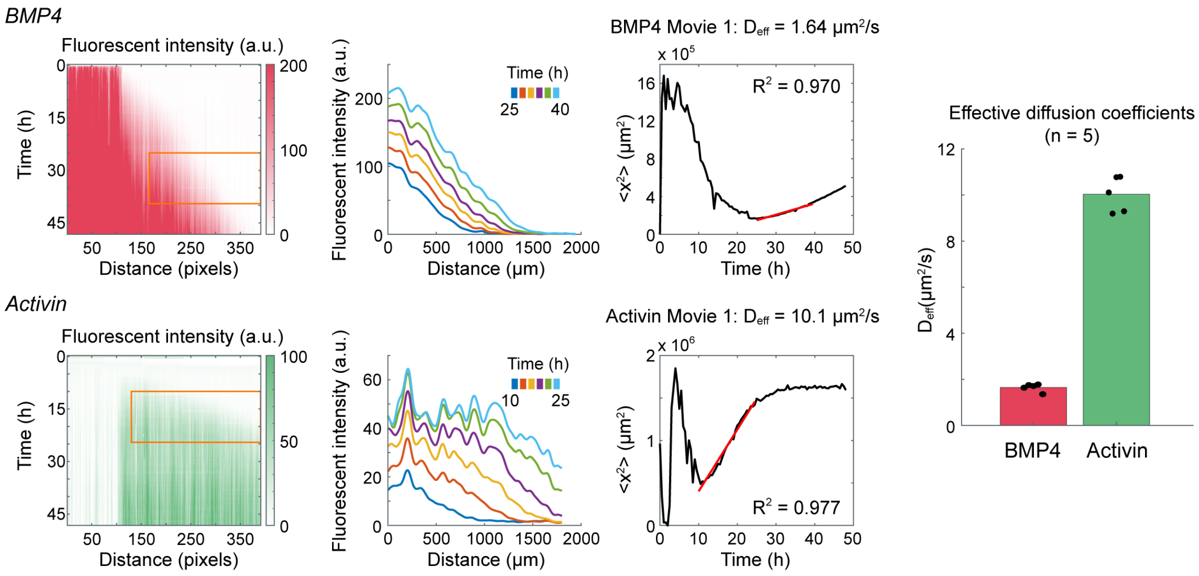


**Fig. S2. Quantification of effective signaling spread from time-lapse reporter movies. (A)** Representative kymographs of BMP4- and Activin-dependent reporter activity from a single movie. Orange boxes indicate the fixed distal receiver-only analysis windows used for moment analysis. **(B)** Representative one-dimensional reporter fluorescence profiles extracted from the movie shown in A at the indicated time points. Profiles were generated by averaging fluorescence intensity along the $y$-axis within the analysis window. **(C)** Reporter-weighted second spatial moment, $M_2(t)$, calculated from the profiles in B. Red lines indicate linear fits over the selected approximately linear intervals, from which the effective signaling-spread coefficient, $D_{\mathrm{eff}}$, was estimated. Panels A to C show the analysis of one representative movie. **(D)** Summary of $D_{\mathrm{eff}}$ estimates from five independent movies for BMP4 and Activin. Bars show the mean, error bars show the sample standard deviation, and individual points indicate values from individual movies.

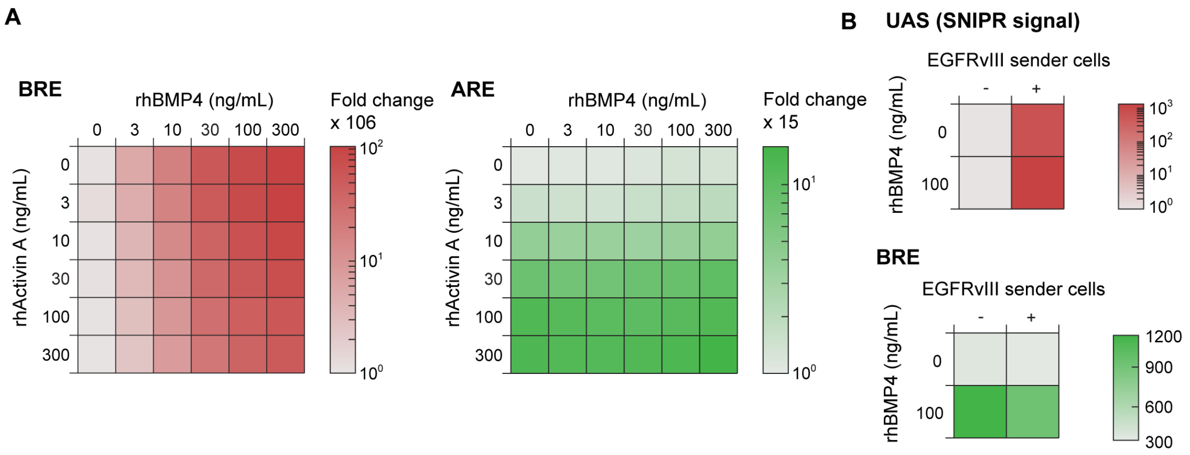


**Fig. S3. Input selectivity and compatibility of BMP4/BRE, Activin/ARE, and SNIPR/UAS signaling modules. (A)** Responses of BRE-driven mCherry (left) and ARE-driven mCitrine (right) reporters in dual-reporter cells across a matrix of recombinant BMP4 and Activin A concentrations. Colors indicate reporter fold change relative to the unstimulated condition, as shown by the corresponding color bars. BRE output varied predominantly with BMP4 concentration, whereas ARE output varied predominantly with Activin A concentration. **(B)** Compatibility of the SNIPR/UAS and BMP4/BRE modules in the Circuit 2 cellular context. Top, UAS reporter output in receiver cells co-cultured with or without EGFRvIII-expressing sender cells, in the absence or presence of 100 ng/mL recombinant BMP4. Bottom, BRE reporter output measured under the same factorial conditions. SNIPR/UAS output was induced by EGFRvIII-expressing sender cells, whereas BMP4 increased BRE output in both sender-cell conditions. Colors indicate reporter fluorescence, with scales shown at right.

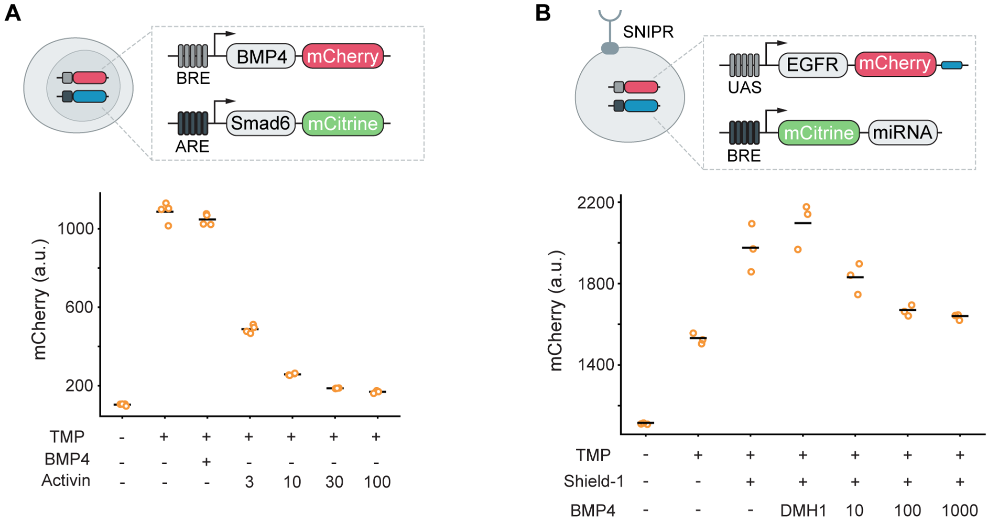


**Fig. S4. Open-loop validation of LALI regulatory modules. (A)** Validation of the Circuit 1 BMP4 self-activation and Activin-dependent inhibitory branches. The diagram shows the BRE-driven BMP4–mCherry module and the ARE-driven Smad6–mCitrine module. mCherry fluorescence was low without TMP, increased following TMP addition, and was further elevated by recombinant BMP4. Under TMP-containing conditions, recombinant Activin reduced mCherry output over the indicated concentration series, consistent with Activin-induced Smad6 suppressing BMP-responsive output. Individual points indicate replicate measurements; horizontal bars indicate means. **(B)** Validation of the Circuit 2 SNIPR self-activation and BMP4-dependent inhibitory branches. The diagram shows the UAS-driven EGFRvIII–mCherry module downstream of SNIPR and the BRE-driven mCitrine–microRNA module. Shield1 increased mCherry output by stabilizing SNIPR. In the presence of TMP and Shield1, DMH1 increased mCherry output, whereas recombinant BMP4 at the indicated concentrations reduced it, consistent with BMP4 signaling inhibiting the SNIPR activation branch through the synthetic microRNA module. Individual points indicate replicate measurements; horizontal bars indicate means.

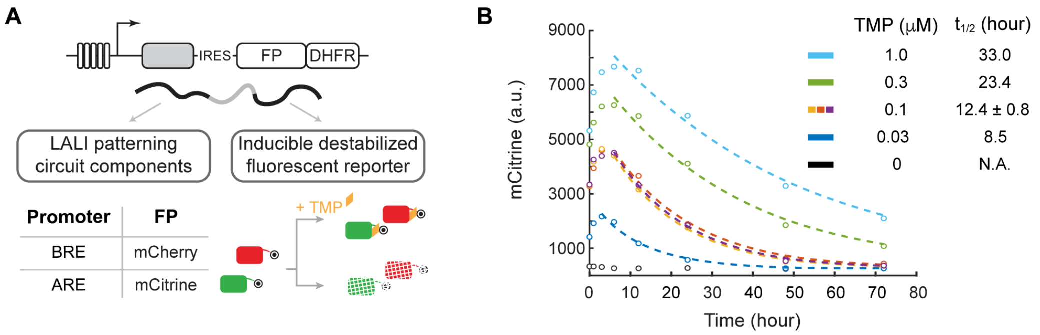


**Fig. S5. TMP-tunable destabilized fluorescent reporters for live imaging of circuit activity. (A)** Reporter design. Each fluorescent protein reporter is encoded on the same transcript as the corresponding LALI circuit component and is translated independently through an internal ribosome entry site (IRES). The reporter is fused to a DHFR destabilization domain, allowing its stability to be regulated by trimethoprim (TMP). BRE-driven mCherry and ARE-driven mCitrine reporters were used to monitor BMP- and Activin-responsive circuit activity, respectively. **(B)** mCitrine fluorescence decay trajectories measured at the indicated TMP concentrations. Dashed lines show exponential fits used to estimate apparent reporter half-lives, which increased with TMP concentration. The table lists the fitted half-life for each TMP condition.

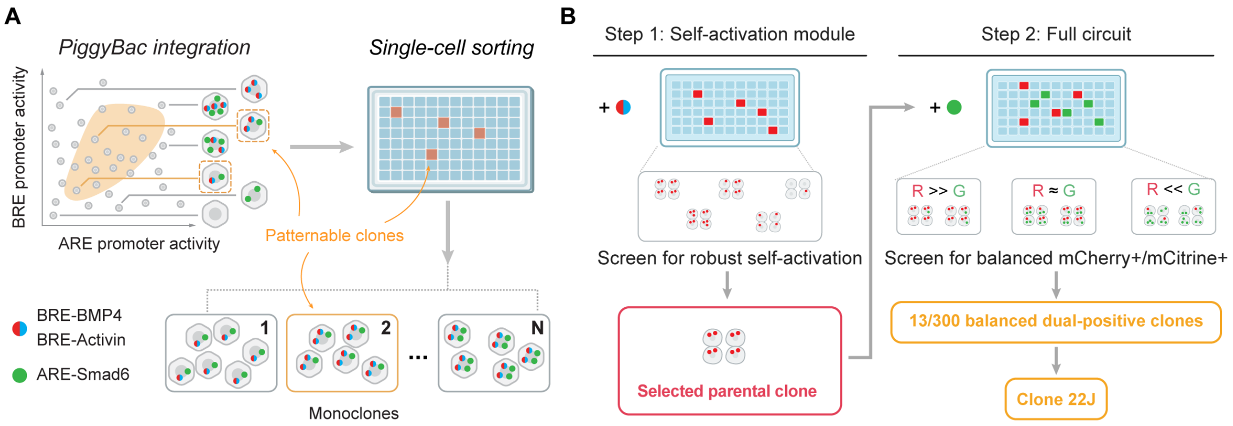


**Fig. S6. Clone-based construction and selection strategy for the BMP4/Activin patterning circuit. (A)** Conceptual scheme of clone selection after piggyBac-mediated integration. Random integration generates monoclonal lines with different effective activities of the BRE- and ARE-driven circuit modules. Single-cell sorting isolates individual monoclonal lines, enabling selection of candidate clones with expression configurations compatible with pattern formation. **(B)** Sequential construction and screening workflow. In step 1, the BRE-driven BMP4 self-activation module was integrated and monoclonal lines were screened for robust self-activation to identify a parental clone. In step 2, the ARE-driven Smad6 inhibitory module was introduced into this parental clone. Resulting monoclonal lines were screened for balanced mCherry and mCitrine reporter expression under reporter-stabilizing conditions. Thirteen of 300 screened clones met this dual-positive criterion and were evaluated in standardized patterning assays; clone 22J was selected for detailed characterization.

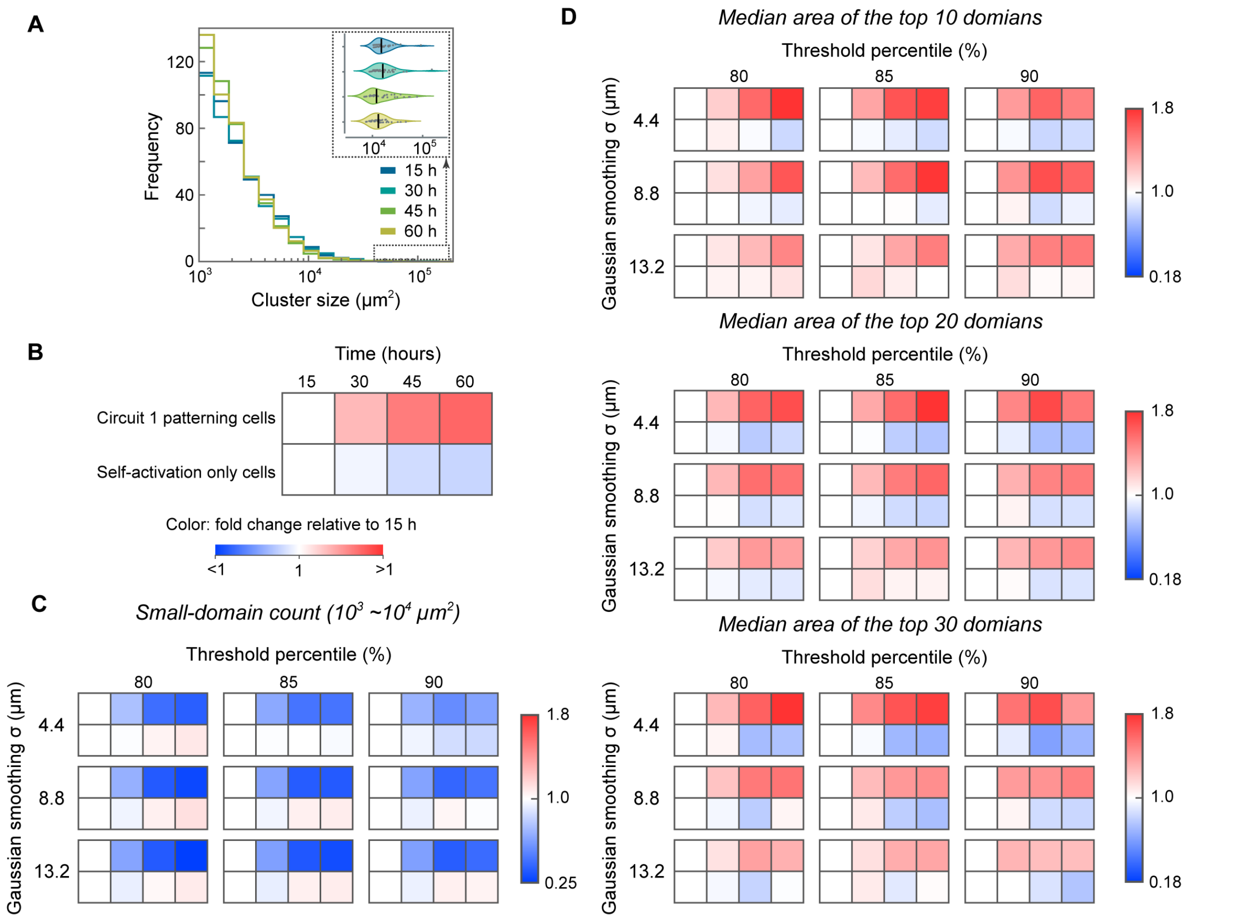


**Fig. S7. Replicate quantification and robustness analysis of Circuit 1 domain competition.** (A) Cluster-size distribution of the top 10% brightest BRE-reporter pixels in the self-activation-only control, in which Activin signaling was fully inhibited by saturating A83-01. Unlike Circuit 1 patterning cells, this control did not show a progressive loss of small domains or enrichment of large domains over time. Inset shows the distribution of the largest-domain areas at each time point. (B) Fold-change summary comparing Circuit 1 patterning cells and self-activation-only control cells over time. Values were normalized to the corresponding 15-h value for each condition. Circuit 1 patterning cells showed a decrease in small-domain abundance and growth of large domains, whereas self-activation-only controls showed little systematic change. (C) Robustness analysis of small-domain counts across image-processing and segmentation parameters. Small domains were defined as connected bright-pixel clusters with areas between $10^3$ and $10^4$ $\mu m^2$. Heatmaps show fold change relative to 15 h across threshold percentiles and Gaussian smoothing widths. (D) Robustness analysis of large-domain growth across image-processing and segmentation parameters. Heatmaps show the median area of the top 10, top 20, or top 30 largest domains, normalized to the corresponding 15-h value. Across a broad range of thresholds and smoothing parameters, Circuit 1 patterning cells showed progressive enrichment of larger domains, whereas the self-activation-only control showed weaker or no consistent change.

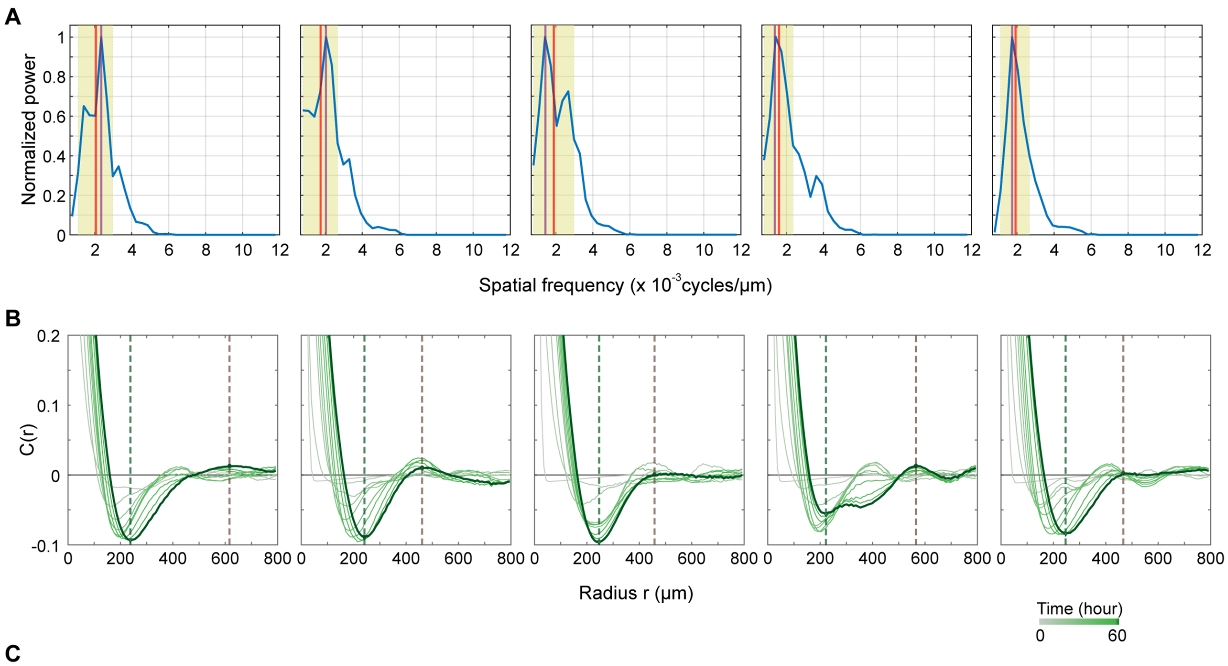


C

Pattern spacing estimation (μm)

| Method | Circuit 1 (n = 5) | Figure reference |
|---|---|---|
| FFT max bin | 582.0 ± 127.4 | Purple solid line in panel A |
| FFT centroid | 546.8 ± 54.9 | Red solid line in panel A |
| C(r) 2x min | 475.2 ± 21.6 | Green dash line in panel B |
| C(r) first peak | 512.2 ± 73.2 | Brown dash line in panel B |

**Fig. S8. Replicate pattern-spacing estimation for Circuit 1.** (A) Radially averaged two-dimensional FFT spectra from five independent Circuit 1 movies after background and uneven-illumination correction. For each movie, normalized power is plotted as a function of spatial frequency. The purple line marks the maximum-power bin, and the red line marks the centroid of the dominant spectral peak. (B) Radially averaged autocorrelation curves, $C(r)$, for the same five independent Circuit 1 movies. Curves are shown over time from 0 to 60 h. Dashed lines mark spacing estimates derived from the first negative minimum and the first weak off-center positive peak. The curves show a reproducible short-range negative correlation but only weak off-center positive peaks, indicating local spatial organization with limited long-range order. (C) Summary of Circuit 1 pattern-spacing estimates from four independent methods. Values are reported as mean ± standard deviation across five movies. The FFT-centroid estimate gives a characteristic pattern spacing of $546.8 \pm 54.9$ $\mu$m. Alternative estimates were $582.0 \pm 127.4$ $\mu$m by FFT max bin, $475.2 \pm 21.6$ $\mu$m by $2\times$ the first $C(r)$ minimum, and $512.2 \pm 73.2$ $\mu$m by the first off-center $C(r)$ peak.

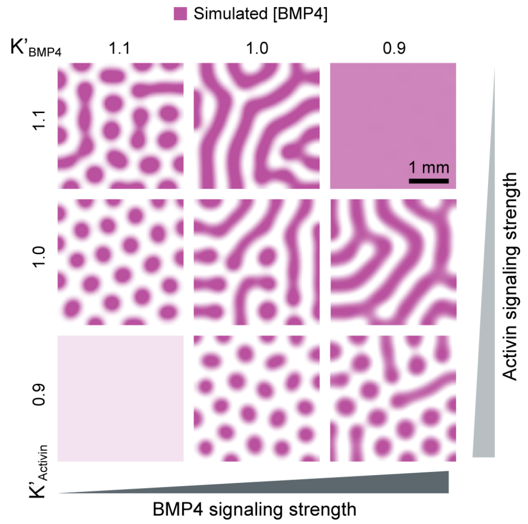


**Fig. S9. Noise-free simulations of Circuit 1.** Simulations were performed under the same conditions as in Fig. 4A, except that stochastic noise was omitted.

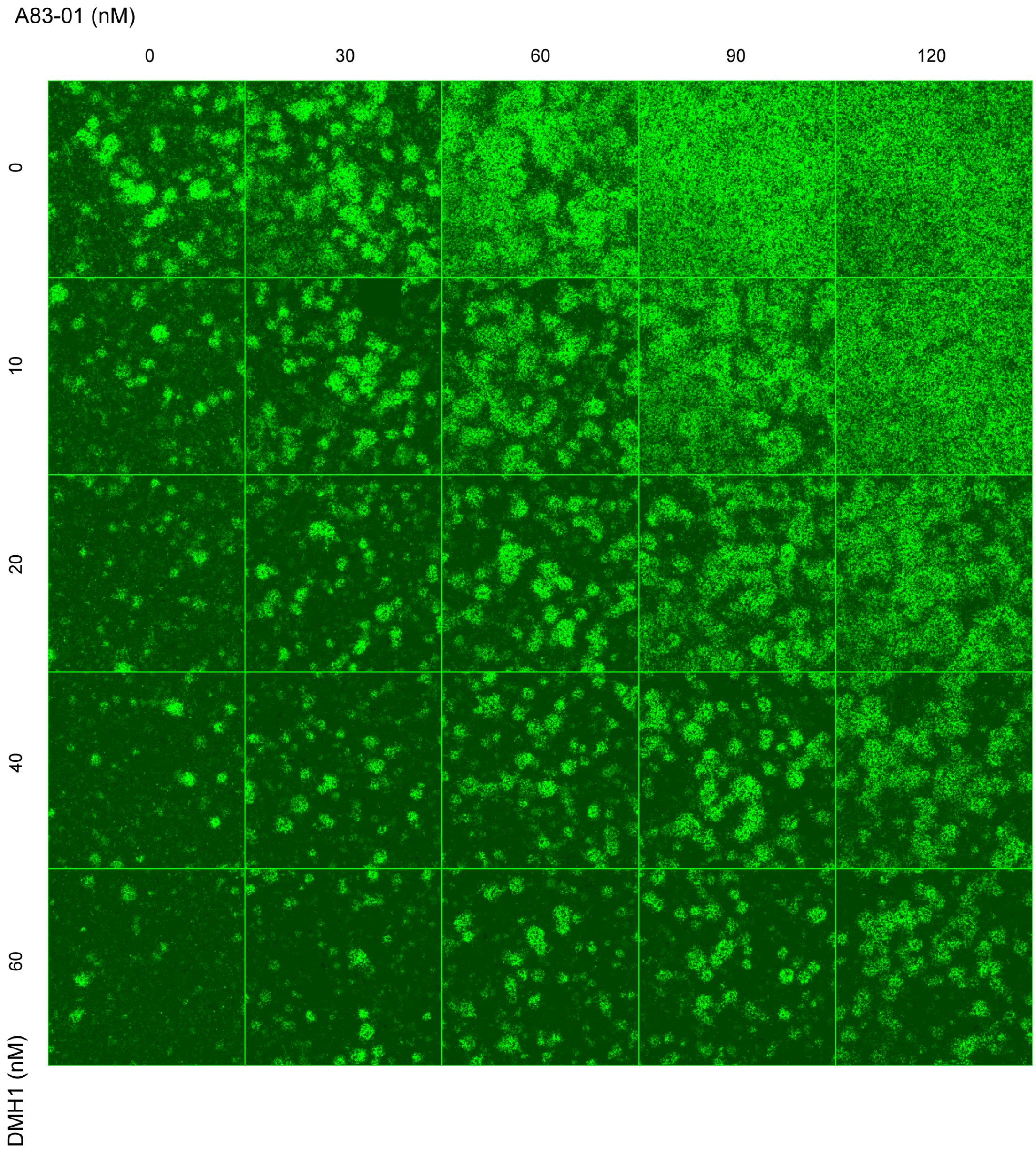


**Fig. S10. Expanded repeat of Circuit 1 DMH1/A83-01 titration.** Representative reporter images from an expanded Circuit 1 titration across DMH1 and A83-01 concentrations. Columns indicate A83-01 concentration and rows indicate DMH1 concentration. This independent repeat reproduced the trend shown in Fig. 4B: reducing Activin-pathway inhibition or altering BMP-pathway activity systematically changed active-domain density and morphology, shifting the pattern between sparse domains and more connected, sponge-like structures.

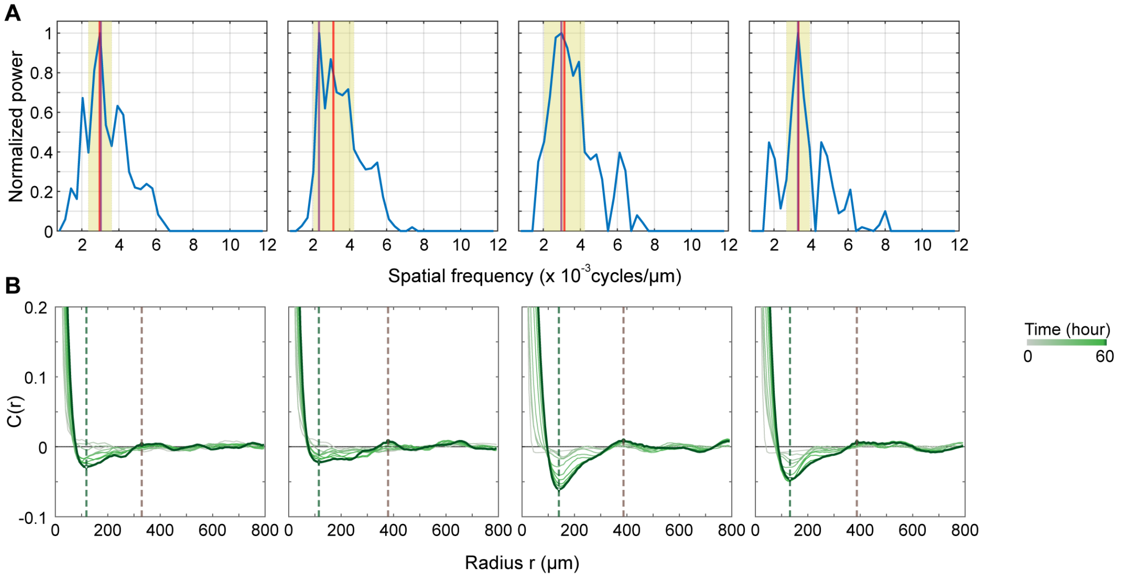


C

Pattern spacing estimation (μm)

| Method | Circuit 2 (n = 4) | Figure reference |
|---|---|---|
| FFT max bin | 349.7 ± 52.2 | Purple solid line in panel A |
| FFT centroid | 319.3 ± 15.3 | Red solid line in panel A |
| C(r) 2x min | 250.8 ± 26.4 | Green dash line in panel B |
| C(r) first peak | 369.6 ± 29.6 | Brown dash line in panel B |

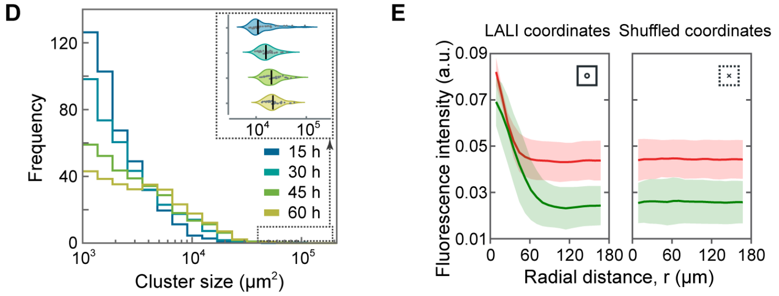


**Fig. S11. Replicate pattern quantification for Circuit 2.** (A) Radially averaged two-dimensional FFT spectra from four independent Circuit 2 movies after background and uneven-illumination correction. For each movie, normalized power is plotted as a function of spatial frequency. The purple line marks the maximum-power bin, and the red line marks the centroid of the dominant spectral peak. (B) Radially averaged autocorrelation curves, $C(r)$, from the same four independent Circuit 2 movies over time. Dashed lines mark spacing estimates from $2\times$ the first negative minimum and the first weak off-center positive peak. Similar to Circuit 1, Circuit 2 showed local spatial organization but weak long-range order. (C) Summary of Circuit 2 pattern-spacing estimates from four methods. Values are reported as mean $\pm$ standard deviation across four movies. The FFT-centroid estimate gives a characteristic pattern spacing of $319.3 \pm 15.3$ $\mu$m. Alternative estimates were $349.7 \pm 52.2$ $\mu$m by FFT max bin, $250.8 \pm 26.4$ $\mu$m by $2\times$ the first $C(r)$ minimum, and $369.6 \pm 29.6$ $\mu$m by the first off-center $C(r)$ peak. (D) Cluster-size distribution of active domains in Circuit 2 over time, quantified from the top 10% brightest pixels. The distribution shifted over time as active domains emerged and reorganized. (E) Event-based superposition analysis of Circuit 2 active domains. Coordinates of active SNIPR/UAS domains were identified and used to align local reporter fields. Shuffled coordinates served as a null control. The aligned profiles reveal local enrichment around active-domain centers, whereas shuffled coordinates produce flat profiles.

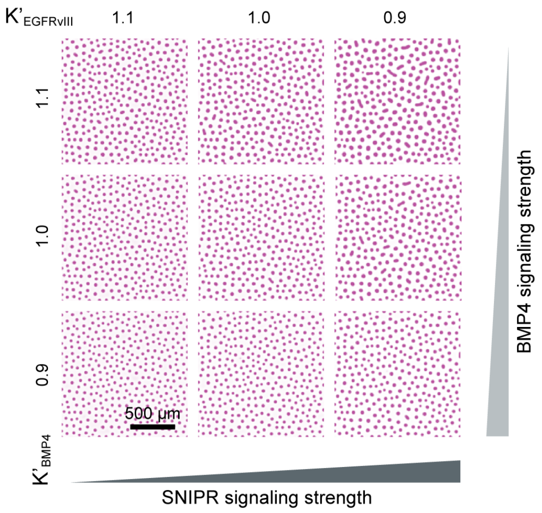


**Fig. S12. Noise-free simulations of Circuit 2.** Simulations were performed under the same conditions as in Fig. 5C, except that stochastic noise was omitted.

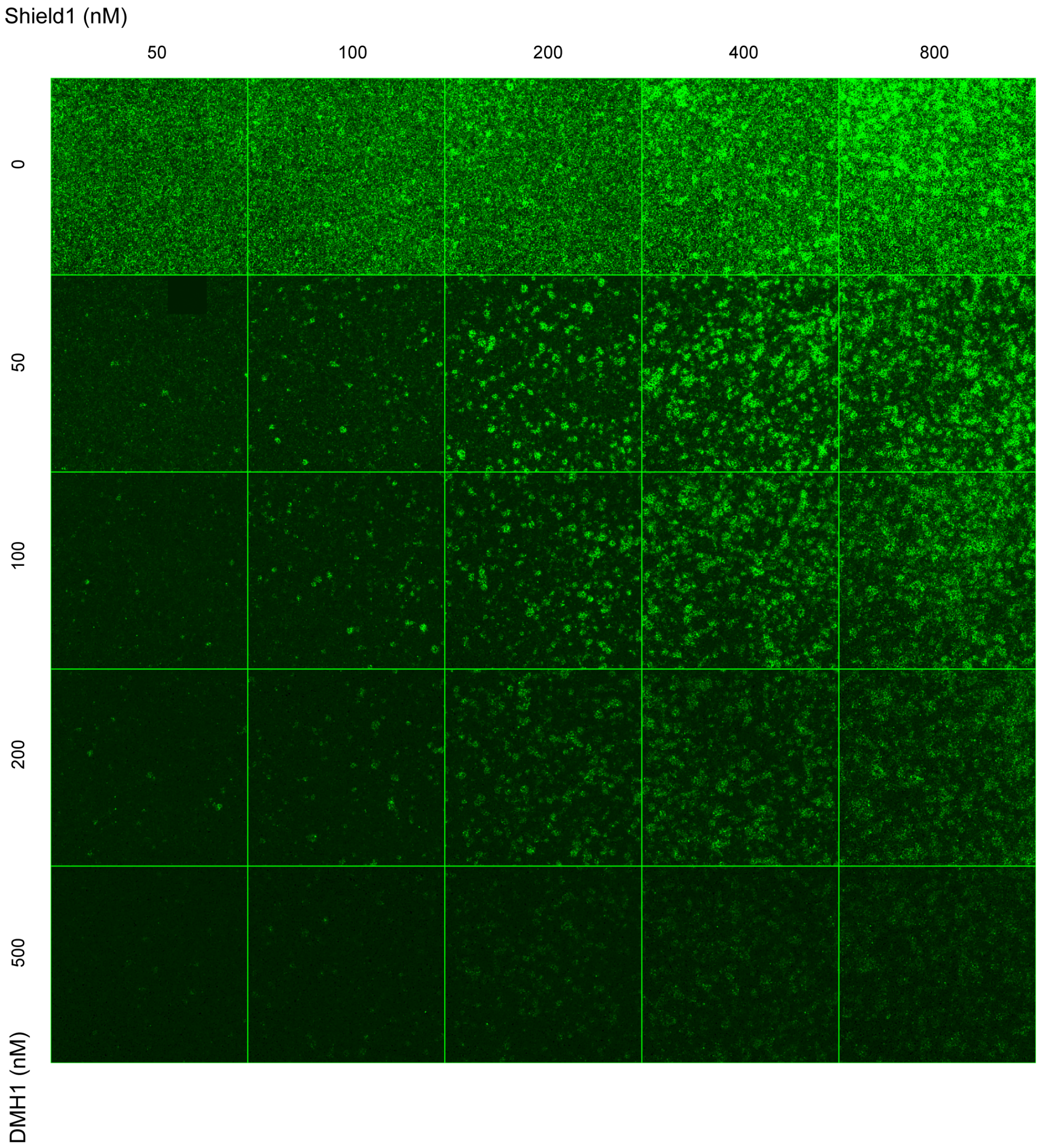


**Fig. S13. Expanded repeat of Circuit 2 Shield1/DMH1 titration.** Representative reporter images from an expanded Circuit 2 titration across Shield1 and DMH1 concentrations. Columns indicate Shield1 concentration and rows indicate DMH1 concentration. This repeat reproduced the trend shown in Fig. 5D: increasing SNIPR stabilization and modulating BMP-pathway inhibition systematically altered Circuit 2 pattern density and reporter activity, supporting the tunability of the SNIPR/BMP4 patterning circuit.

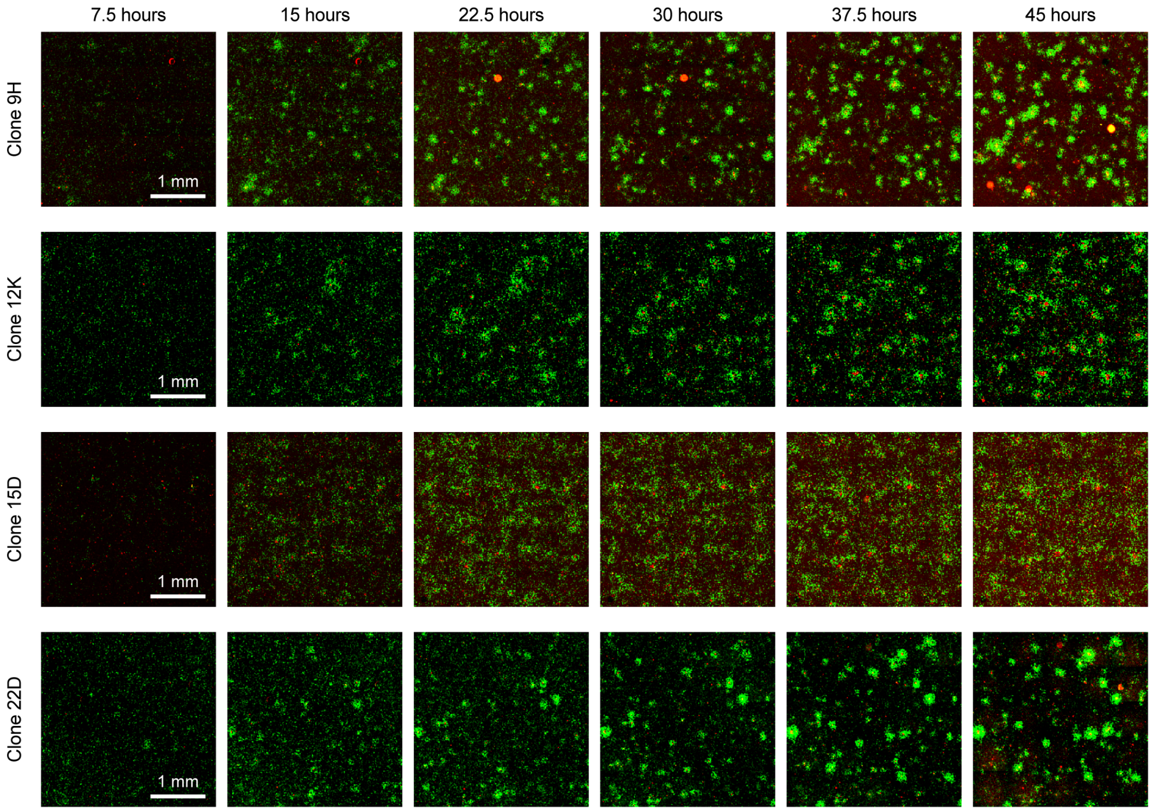


**Fig. S14. Time-lapse dynamics of additional independently derived BMP4/Activin circuit clones.** Representative time-lapse images of four full-circuit monoclonal lines under the standardized patterning condition. Columns indicate the time after pattern initiation. BRE-driven mCherry is shown in red, and ARE-driven mCitrine is shown in green. Clone 9H and clone 22D developed dynamic localized reporter-positive domains, whereas clone 12K and clone 15D showed broader spatial heterogeneity with less clearly resolved domain organization. Scale bars, 1 mm.

### Supplementary Movie Legends

#### Movie S1. Time-lapse fluorescence imaging of the RD condition

Representative time-lapse imaging of the Circuit 1 RD condition (TRP734-Well005), showing merged mCherry (BRE; red) and mCitrine (ARE; green) fluorescence over 60 h. Images were acquired every 1.5 h. A fixed intensity range was applied to each fluorescence channel throughout the movie. Time is indicated in hours. Scale bar, 500 $\mu$m.

#### Movie S2. Time-lapse fluorescence imaging of the control condition

Representative time-lapse imaging of the Circuit 1 control condition (TRP776-Well006), showing merged mCherry (BRE; red) and mCitrine (ARE; green) fluorescence over 60 h. Images were acquired every 1.5 h. The same fixed channel-specific intensity ranges used for Movie S1 were applied throughout the movie. Time is indicated in hours. Scale bar, 500 $\mu$m.

#### Movie S3. Higher-magnification comparison of the RD and control conditions

Side-by-side time-lapse comparison of mCherry (BRE) fluorescence in the RD (left; TRP734, Well005) and control (right; TRP776, Well006) conditions over 60 h. Each panel shows the same-sized 300 $\times$ 300-pixel region selected from the corresponding field of view. Images were acquired every 1.5 h and displayed with the Reds color map. In Movie S3.mp4, each condition is displayed using its own color scale, normalized to the minimum and maximum fluorescence intensities across the entire movie for that condition. Time is indicated in hours. Scale bar, 100 $\mu$m.

#### Movie S4. Time-lapse comparison of selected competition regions in the RD and control conditions

Side-by-side time-lapse comparison of mCherry (BRE) fluorescence within matched, rotated competition-region crops from the RD (left; TRP734, Well005) and control (right; TRP776, Well006) conditions over 60 h. Each crop spans 240 $\times$ 120 pixels. Images were acquired every 1.5 h and displayed with the Reds color map. In Movie S4.mp4, each condition is displayed using its own color scale, normalized to the minimum and maximum fluorescence intensities across the entire movie for that condition. Time is indicated in hours. Scale bar, 100 $\mu$m.

#### Movie S5. Pattern-formation dynamics of TRP808 Circuit 1 under 25 induction conditions

Pattern-formation dynamics of TRP808 Circuit 1 under 25 induction conditions, presented as a $5 \times 5$ montage of mCitrine (ARE) fluorescence. Images were acquired every 3 h, and the displayed sequence spans 0–60 h. Time is indicated in hours. Scale bar, 1 mm.

#### Movie S6. Time-lapse pattern formation of TRP833 Circuit 2

Time-lapse fluorescence imaging of Circuit 2 (TRP833-Well002) under the patterning condition shown in Fig. 5A. Merged mCherry (UAS; red) and mCitrine (BRE; green) fluorescence is shown over 60 h. Images were acquired every 1.5 h, and fixed intensity ranges were applied to each fluorescence channel throughout the movie. Time is indicated in hours. Scale bar, 500 $\mu$m.

#### Movie S7. Pattern-formation dynamics of TRP830 Circuit 2 under 25 induction conditions

Pattern-formation dynamics of TRP830 Circuit 2 under 25 induction conditions, presented as a $5 \times 5$ montage of mCitrine (BRE) fluorescence. Images were acquired every 3 h, and the displayed sequence spans 0–60 h. Time is indicated in hours. Scale bar, 1 mm.

#### Movie S8. Time-lapse comparison of mCherry and mCitrine fluorescence in four clones

Time-lapse fluorescence imaging of four Circuit 1 clones, shown from left to right as clone 9H, clone 12K, clone 15D, and clone 22D. mCherry (BRE) and mCitrine (ARE) fluorescence are shown in red and green, respectively. Images were acquired every 1.5 h, and the displayed sequence spans 1.5–46.5 h. For each clone, the red and green channels share a single fixed intensity range that is applied throughout the movie, preserving within-clone red-to-green intensity relationships over time. Time is indicated in hours. Scale bar, 1 mm.
